# Replication Timing Uncovers a Two-Compartment Nuclear Architecture of Interphase Euchromatin in Maize

**DOI:** 10.1101/2025.07.11.664491

**Authors:** HS. Akram, E.E. Wear, L. Mickelson-Young, Z.M. Turpin, L. Hanley-Bowdoin, W.F. Thompson, L. Concia, H.W. Bass

## Abstract

Genome replication is temporally regulated during S phase, with specific genomic regions replicating at defined times in a process that is known as replication timing (RT). Based on 3D cytology in replicating nuclei, we previously proposed a “mini-domain chromatin fiber RT model” for maize euchromatin that suggested it is subdivided into early-S and middle-S compartments distinguished by chromatin condensation and RT. However, whether this compartmentalization reflects a general nuclear architecture that persists throughout the cell cycle was unclear. To test this model, we conducted two orthogonal assays – Hi-C for genome-wide interaction data and 3D FISH for direct visualization of chromatin organization. Hi-C eigenvalues and insulation scores revealed distinct patterns of early-S regions having negative insulation scores with long-range contacts while middle-S regions showed the opposite. Early-S regions correlated more strongly with epigenomic signatures of open, transcriptionally active chromatin than middle-S regions. 3D oligo FISH painting confirmed that early-S and middle-S regions occupy adjacent but largely non-overlapping nucleoplasmic spaces during all interphase stages, including G1. Our findings redefine the maize euchromatin “A” compartment as having two distinct, closely interspersed subcompartments, early-S and middle-S, underscoring the importance of replication timing (RT) as a defining feature of chromatin architecture and genome organization.

## INTRODUCTION

The eukaryotic cell cycle consists of a widely conserved series of events, including cell growth, DNA replication, and division into two daughter cells. DNA replication occurs during the S phase, and the specific time within S phase when each genomic region replicates can be measured and annotated as replication timing (RT) ^1–3^. The fact that some cells or tissues have different RT profiles reveals the existence of an underlying RT program coupled to development or differentiation ^4,5^. The RT program is generally recognized as ensuring the faithful reproduction and transmission not only of the nucleotide sequence but also of the chromatin state, contributing to the propagation of epigenomic cell type identity and functions ^6^.

Most eukaryotic RT knowledge comes from yeast and mammals, where complex temporal programs have been described and extensively studied ^7,8^. However, the regulatory mechanisms that govern temporal RT programs remain understudied in plants ^9^. Notably, plants lack homologs of key RT regulators such as geminin and Rif1, which are essential for coordinating the cell cycle and RT in yeast and mammals (reviewed by ^10,11^). Similarly, plants and mammals exhibit some key differences in their chromatin organization as characterized by proximity ligation assays. A prominent feature of mammalian chromatin architecture is the presence of topologically associating domains (TADs), which partition the genome into domains spanning hundreds of kilobases ^12^. These TAD boundaries are typically enriched for architectural proteins such as CTCF and cohesin. In contrast, plants are reported to lack any clear CTCF homolog and cohesin-enriched TAD boundaries ^13–15^. Together, these observations strongly suggest that there are fundamental differences in the higher-order organization of plant chromatin. These differences along with the vast evolutionary distance between plants and opisthokonts highlight the importance of studying DNA RT in plant models.

Early studies of plants concluded that genome replication is essentially biphasic, with early-replicating and late-replicating regions ^16,17^. Later studies greatly improved the resolution of the RT profile by using nucleotide analogs such as 5-bromo-2′-deoxyuridine (BrdU) ^18^ and 5-ethynyl-2’-deoxyuridine (EdU) for pulse-labeling replicating DNA. EdU has become the analog of choice because it does not require acid or heat denaturation for detection and, instead, is directly conjugated with click chemistry to add a fluorescent dye used for both fluorescence activated nuclei sorting (FANS) and microscopy ^19^. These EdU-labeled nuclei can be separated by flow cytometry from unlabeled G1 and G2 nuclei and further divided by increasing DNA content into subpopulations representing sequential stages of S phase. Three separated populations, early-S, middle-S, and late-S, have been used for microscopic analysis and sequencing of replicated DNA (Repli-seq) ^20^. The first genome-wide RT maps in plants were produced for *Arabidopsis thaliana* and maize (*Zea mays* L.) using these techniques ^2,21^. The comparative analysis of early-S, middle-S and late-S replication in Arabidopsis revealed that early-S and middle-S are quite similar to each other and distinct from late-S. Early and middle replicating domains are enriched with open chromatin as defined by nuclease sensitivity and histone acetylation markers with early-replicating regions being gene-rich, while late-replicating domains are characterized by an enrichment of transposons and repressive epigenetic markers ^13,18,21,22^. Although *Arabidopsis* has provided a foundational understanding of plant RT, its small genome size and high gene density are unusual among plant genomes. Plants with larger genome sizes, such as maize (2*n*=2X=20) with a 1C value of 2.3 Gb, have expanded intergenic content, which generally tends to replicate primarily during middle-S and exists in a slightly more compacted state compared to the early-S replicating chromatin ^13,23,24^.

The maize genome, which is about two thirds the size of the human genome and contains a complex array of transposons, is more typical of higher plants. This fact, together with the fact that maize has been the subject of detailed genetic and cytogenetic studies for more than 100 years ^25,26^, makes it an attractive system for exploring the control of replication in plants with complex genomes and chromatin architecture. Furthermore, as one of the world’s most agriculturally significant crops, insights into the replication process in maize are relevant not only for fundamental research but also for strategies to enhance crop performance and resilience.

Our group has developed a powerful experimental system to characterize replication in actively growing, intact maize root tip meristems. This system has provided key insights and enables examination of replication patterns, genome features, and associated chromatin structure parameters at different stages of the cell cycle ^20,27^. The maize root tip system examines nuclei from cells growing in their native, organismal, and developmental context, unlike nearly all of the mammalian models that depend primarily on cell culture systems. Although heterochromatic regions in maize replicate late, and many genes replicate early as they do in most other eukaryotic systems, the spatiotemporal patterns of early and middle S-phase replication in maize differ significantly from those in mammalian and yeast systems. In mammals, DNA synthesis occurs at different nuclear regions throughout S phase, classified as 5 sequential stages referred to as patterns 1-5 ^28^ or Types I-V ^29^, in which the first, third, and fifth stages represent the earliest, the middle most, and latest stages of S phase, respectively. Despite the different naming conventions, there is agreement on the nuclear distribution of replication in mammalian nuclei, which starts with broad distribution in euchromatic portions of the nucleoplasm at early S phase, then in perinuclear and perinucleolar regions during middle S phase, followed by heterochromatic portions of the nucleoplasm at late S phase ^6,28–31^. Many mammalian RT studies compare early to late RT, and genomic regions where they switch, with relatively few studies focusing directly on middle-S ^4,32,33^. In maize, DNA synthesis is thoroughly dispersed throughout the nucleus during both early-S and middle-S phase and becomes more punctate and clustered only in late-S phase. Analysis of DNA RT in maize shows that early-S phase replicative labeling primarily colocalizes with regions of relatively weak 4’,6-diamidino-2-phenylindole (DAPI) fluorescence, indicating a low DNA density, while middle-S phase EdU labeling aligns with regions of euchromatin showing stronger DAPI staining. Based on these findings, we proposed a "mini-domain chromatin fiber RT model”, which states that maize euchromatin exists as a closely interspersed mixture of two subcompartments, each characterized by distinct RT and chromatin morphology ^24^.

To test the mini-domain model, we employed a combination of proximity ligation (Hi-C) and quantitative in situ hybridization (3D-FISH) techniques in this study. Here we provide new evidence supporting this model of maize euchromatin, showing that the described relationships hold throughout the cell cycle. Our results support a view of the plant euchromatin global "A" compartment as largely composed of two spatially and epigenetically distinct, but closely interspersed, compartments.

## METHODS

### Plant materials

Experiments were done using maize inbred B73 root tips or earshoots. For root tips, as described in a previous protocol ^34^, seeds were surface sterilized and imbibed overnight with constant stirring and aeration. Seeds were surface-sterilized again with 0.5% sodium hypochlorite, 0.05% Tween-20 solution, and then rinsed with water. Seeds were germinated in boxes with paper towels moistened with sterile water at 28°C for 3 days under continuous light (Feit Electric OneSync LED light system, 3000K, 300-400 lux). The material for biological replicates were grown independently and harvested on different days. For earshoots, field-grown maize plants were harvested between 9-11am on sunny days, and earshoots ranging in size from 0.5-1 cm were flash frozen in liquid nitrogen, and pooled (15-20 earshoots) for storage at -80°C as described in previous paper ^35^.

### Repli-seq realigned to B73v5

For this study, we used previously published Repli-seq data generated from maize B73 root tips that were EdU labeled, dissected (terminal 1-mm) and fixed as described ^2^. In isolated nuclei, the EdU was conjugated to Alexa Fluor 488 (AF-488), and total DNA stained with DAPI for flow sorting of early-S, middle-S, and late-S nuclei, as well as non-replicating G1 nuclei, based on both EdU incorporation and DNA content (like in Figure 1a). After DNA extraction, the EdU labeled DNA was immunoprecipitated from the S-phase fractions and sequenced. The raw sequence data from early-S, middle-S, or late-S and the G1 reference libraries were aligned to the current working version of the maize genome, Zm-B73-REFERENCE-NAM-5.0 (B73v5), using BWA mem (v0.7.15) ^36^ with default parameters. Aligned sequences that were properly paired and uniquely-aligned (MAPQ score >10), were retained with SAMtools ^37^ and duplicates removed using Picard MarkDuplicates (v2.7.1). For genome segmentation and annotation, we used the previously custom-built Repliscan software as described ^38^. Highly correlated replicates ^2^ were pooled to produce read densities in different bin sizes (1, 3, 5 and 10 kb) across the genome. The various bin sizes were used for resolution analyses and all four bin sizes for all three RT samples are available as bigwig files. Repliscan normalization was achieved in two ways; (1) by sequence depth scaling to 1x coverage using the reads per genomic content (RPGC) method, and (2) by normalization to a non-replicating reference (G1) to correct for sequencing biases, read mappability variation, and copy number. Genomic bins with the lowest 1% from a reads per bin distribution in G1 were also removed. The final replication profile data for each of the S-phase samples was calculated as a ratio of the S-phase sample to the G1 reference value in each bin, and represents an estimate of the intensity of replication activity in that bin (“replication signal”) for that portion of S phase. Finally, replication signal profiles for each S-phase sample were Haar wavelet smoothed ^39^. The final step in the Repliscan segmentation incorporates the replication signal for each S-phase portion in each bin and makes a qualitative call of the predominant replication time in that bin. The largest replication signal value is always classified as replicating, and the algorithm allows multiple time classifications (e.g., early-mid or middle-late) when another signal is within 50% of the highest value. The parameters for maize B73 described in Mickelson-Young *et al.,* (2022) were used except for the following minor modification: each bin was assigned to a RT segment class using a segmentation threshold of 1.0 (--*threshold* value, --*value* 1.0), so that the segmentation algorithm included any window with early, middle or late replication signals that are greater than the G1 reference signal (> ratio of 1.0). The resulting Repliscan segmentation file assigned unique and mutually exclusive RT classes across the genome. We primarily used the early (E), middle (M), or late (L) segments, which accounted for ∼30%, ∼35%, or ∼24% of the maize genome, respectively.

**Fig. 1.**
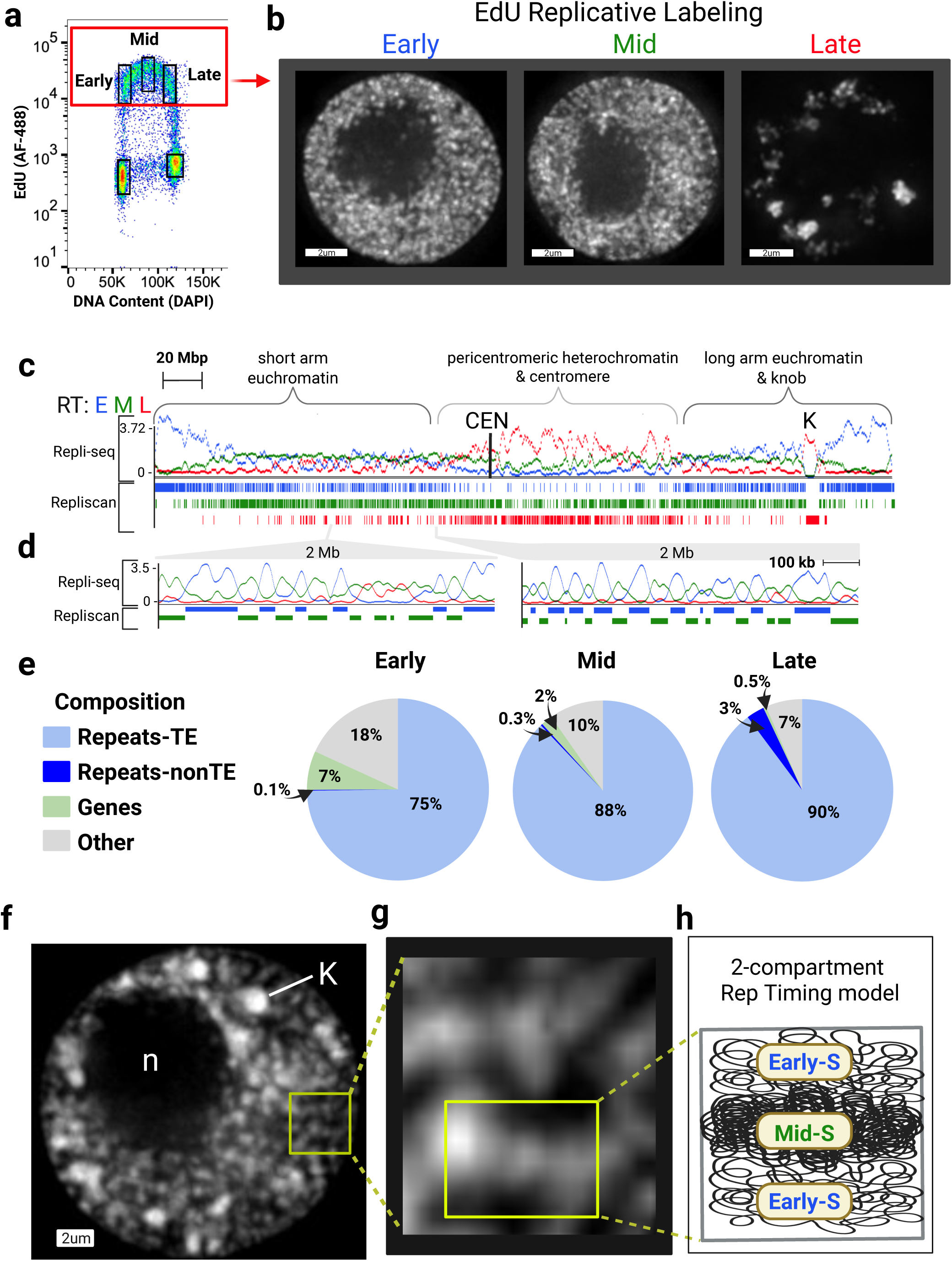
Repli-seq scheme and mini-domain chromatin fiber RT model. **(a)** DAPI and EdU stained nuclei were analyzed via flow cytometry using 355 nm (UV) and 488 nm (blue) lasers. The bivariate plot displays DNA content (DAPI fluorescence with emission filter 460 ± 50 nm) and EdU incorporation (AF-488 fluorescence with emission filter 530 ± 40 nm). Rectangles indicate the G1, early-S, middle-S, late-S and G2 nuclei that were sorted **(b)** Cytology of early-S, middle-S and late-S sorted nuclei. Images were collected using 3D deconvolution microscopy, corrected for wavelength-dependent chromatic aberration, and displayed as grayscale. Each nucleus is shown with intensity-averaged projections spanning 2 µm (10 Z-sections) from the center. **(c)** A schematic representation of euchromatin and heterochromatin distributed along chromosome 5, including a centromere (CEN) ^80,81^ and knob (K) on the long arm. The Repli-seq profiles and Repliscan segments spanning the entire chromosome are shown with early RT (blue), middle RT (green), and late RT (red). **(d)** UCSC Genomaize browser view zoomed into two 2-Mb non-heterochromatin regions with Repli-seq profiles and RT segments defined by Repliscan **(e)** The composition of early-S, middle-S and late-S is shown in pie charts with the legend to the left. **(f)** DAPI stained image of a maize interphase nucleus. The nucleolus is labeled “n”, and one of the knobs is highlighted as “K”. The yellow box indicates the euchromatin region magnified in the next panel. **(g)** A zoomed-in area of the euchromatin region with fiber-like structures. **(h)** Illustration depicting two euchromatin sub-compartments: early-S (thin/gray) and middle-S (thick/black).

### Root tip and earshoot nuclei for Hi-C

For Hi-C on seedling root nuclei, the terminal 0-1 mm of the primary and first two seminal root tips were dissected and crosslinked with 1% formaldehyde for 15 min. Nuclei were isolated, stained with DAPI (No EdU), and flow sorted on a BD FACSAria III equipped with UV (355 nm) and blue (488 nm) lasers and using flow cytometer settings and doublet discrimination gating as previously described ^40^. The sorted nuclei were collected in ∼1.8 mL CLB**^+^** buffer (cell lysis buffer with 1mL DTT and protease inhibitor tablet) as described ^34^. After excluding doublets, debris and endocycling nuclei, approximately 1.3-1.5 million nuclei were collected as input for each biological replicate of the Hi-C assay. A representative FANS profile is shown in Supplementary Figure 1c.

For Hi-C on field-grown earshoot nuclei, we used the method of Savadel *et al.,* (2021) ^35^ to isolate nuclei. The frozen tissues were ground in liquid nitrogen to a fine powder, and the nuclei were crosslinked in 1% formaldehyde. The fixation reaction was stopped by addition of 0.1 vol (∼1 mL) of 2.5 M glycine. The tissue was then mechanically disrupted with a Polytron (3 x 10 seconds at ∼ 1/5 maximum speed) to liberate nuclei. Nuclei were pelleted by centrifugation at 2000 x𝙜 for 15 min at 4°C and resuspended in a 15 mL CLB**^+^** buffer. Total earshoot nuclei were used for the Hi-C assay.

### Preparation and sequencing of Hi-C libraries from maize root tip and ear shoot

The Hi-C libraries were generated essentially following the protocol first described for maize leaf by Dong *et al.* (2017)^41^ and later detailed in a methods chapter by Dong and Zhong (2020) ^42^.

We made some minor modifications to accommodate the library kits currently used by the Molecular Cloning Facility (Department of Biological Science, Florida State University, Tallahassee, FL) and reduced washing steps to avoid nuclei loss as described in the wet bench full protocol. Among the key steps were permeabilization of nuclei with 0.5% (w:v) SDS for 5 min at 65°C, DNA digestion with *Dpn-II* restriction enzyme, overhang repair with Klenow and biotinylated nucleotides, and ligation with T4 DNA ligase. Ligated DNA was biotin affinity purified, crosslinks reversed, and sheared to 300-500 bp to yield DNA fragments for library construction using NEBNext Adapters for Illumina. Libraries from both root tips and earshoots were sequenced using Illumina Hi-Seq 2500 paired-end 150-bp chemistry (Translational Science Lab at College of Medicine, Florida State University, Tallahassee, FL) to obtain library sequences at a depth ranging from 127M to 309M (Supplementary Table1) per replicate, with four replicates for root tips and three for earshoots. We used fastqc (v0.12.1) to assess the read quality and then performed trimming using trimmomatic (v0.39) to remove low-quality bases and adapter sequences. The trimmed reads were used in the HiC-pro pipeline as described in the HiC-Pro Data Processing Pipelines file.

### Nuclei isolation for MMOA-seq

For Mononucleosomal MNase-defined Cistrome-Occupancy Analysis (MMOA-seq), nuclei preparation was done using our previously published protocol ^2,20^. Briefly, maize seedling roots were labeled in 25 µM EdU for 20 min, rinsed and placed in 100 µM thymidine to stop labeling. The terminal 1-mm maize root tip segments were excised, fixed with 1% formaldehyde and lysed to isolate nuclei for MMOA-seq. Incorporated EdU was conjugated to AF-488 using a Click-iT EdU Alexa Fluor 488 imaging kit (Life Technologies). The nuclei were then counterstained with DAPI using a cell lysis buffer containing 2 μg/mL DAPI and 40 μg/mL Ribonuclease A and filtered through a 20-μm nylon mesh filter (Partec). Flow sorted G1 nuclei were used for MMOA-seq library preparation.

### MMOA-seq library preparation and Sequencing

MMOA-seq libraries were prepared using nuclei in 500-µL aliquots per replicate. Briefly, 60 µL of nuclei were distributed to each of four 1.5 mL screw cap tubes. 10x MNase working dilutions (25 U/mL, 12.5 U/mL, and 6.25 U/mL) were prepared from a 20,000 U/mL stock. 6.7 µL of each working dilution (or MDB) were added to each of the 60 µL nuclei aliquots and immediately vortexed and briefly centrifuged. Complete digestion reactions were transferred to a 37°C shaking water bath and incubated for 15 min. Reactions were promptly stopped after 15 min by addition of 5 µL of 0.5 M EGTA (pH 8.0). Digested chromatin was then decrosslinked overnight at 65°C in 1% SDS, 150 mM NaCl, 20 µg/mL Proteinase K. DNA was purified by 25:24:1 Phenol:Chloroform:Isoamyl alcohol (pH 8.0). Nucleic acids were precipitated and the pellets were washed with ice cold 70% ethanol and air dried before being redissolved in 100 µL of 10 mM Tris EDTA buffer.

For each library, 75 ng of selected digests (1.25 and 2.5 U/mL MNase) were combined and diluted to 50 µL with 0.1X TE. Sequencing libraries were made according to the NEBNext Ultra II DNA Library Prep Kit for Illumina with the following modifications to the size selection protocol for retention of small fragments (<200 bp): the first bead addition used 50 µL of beads, the second used 100 µL of beads. These size selection steps were chosen to collect both mononucleosome-sized DNA and smaller sub nucleosome-sized DNA fragments for sequencing library preparation. A total of 8 cycles of PCR were completed at the barcoding step.

Libraries were quantified by Qubit dsDNA HS fluorimetry and KAPA qPCR. DNA fragment size distribution was analyzed on an Agilent High Sensitivity D1000 Screentape assay. An equimolar pool of all 24 MMOA-seq libraries was prepared and sequenced using 50-bp paired-end reads on a Novaseq S1 flow cell at the College of Medicine Translational Science Lab, Florida State University, Tallahassee, FL.

### HiC-Pro analysis of Hi-C data

HiC-Pro ^43^ is a custom bioinformatic software for the analysis of Hi-C datasets. The pipeline performs sequential steps, including short read mapping, detection of valid ligation fragments, and various quality controls. The output consists of inter- and intra-chromosomal contact maps at various resolutions.

Trimmed and filtered paired-end reads were aligned to B73v5 maize reference genome using bowtie-build software (v2.4.4). Then HiC-pro (v3.1.0) was used to generate the contact matrices. The genome was divided into non-overlapping bins of equal size at different resolutions (1 kb, 5 kb,10 kb, 50 kb, 250 kb, 500 kb, and 1mb) and contacts were scored in each bin. The frequency of interaction between bins was represented by bi-dimensional heatmaps (“contact matrices’’), containing both inter- and intra-chromosomal contacts. Contact matrices were visualized using Juicebox ^44^, and used to calculate a Pearson correlation matrix for observed/expected intra- and inter-chromosomal interactions. The KR normalization method was applied to all Hi-C matrices. KR is an implicit normalization method through matrix balancing, and is used as a valid alternative to explicit normalization of sequencing biases ^45,46^.

### Insulation score and Eigenvector analysis

To systematically identify folding structures at the local scale, we applied several techniques, including Principal Component Analysis (PCA), the Insulation Index method and Virtual 4C analysis. We ran chromosome-wide PCA to calculate the first eigenvectors (principal components) for three binning resolutions (25 kb, 50 kb, and 100 kb) and processed files will be available on figshare. The eigenvector (EV) values are commonly used to infer genomic compartments as initially described in Lieberman-Aiden *et al.* (2009) ^47^. We also performed chromosomal block-based eigenvector (BEG) analysis at 50-kb resolution, using the constrained hierarchical clustering method as described for maize by Dong *et al.* in 2017 ^41,48^. A series of insulation score analyses ^49,50^ were calculated, quantifying the cumulative frequency of interactions in each bin size series (1 kb, 10 kb, 25 kb, and 50 kb) and visualized within 1-mb or 2.5-mb sliding windows using the bin sizes as step sizes. We calculated the log2 of the observed/expected interaction frequencies using the median scores for the expected values.

Valleys/minima (negative values) indicate loci of reduced frequency of interaction with flanking regions, whereas local maxima (positive values) reveal folding structures.

Virtual 4C analysis ^51^ was centered over “viewpoints” of interest (“baits”), chosen in our case among replication segments of different classes, early-S or middle-S. We scored all the valid interactions between each viewpoint and the rest of the chromosome, and compared the frequency of early to early and middle to middle versus early to middle interactions.

### Chromatin state analysis using a Hidden Markov Model

For inputs used with chromatin state analysis, we obtained published data sets from B73 maize root tip samples, including histone modification ChIP-seq ^2,52^, MNase-seq ^53^, RAMPAGE ^52^ and Repli-seq ^2^. The data source files are cited and the raw fastq files were aligned to B73v5 as described in GitHub.

Briefly, fastQC software (v0.12.1) was used for quality control, and low-quality reads were filtered out. Adapter cleanup was done by trimmomatic (trimmomatic_latest.sif). Bowtie2 software (v2.4.4) was used to align sequencing reads to the reference genome (B73v5 in maize) with the default parameters. Duplicate reads were removed by samtools software (v1.9-4). Bam files were converted into bed files and used as an input for chromHMM binarization.

The Hidden Markov Model was applied to aggregate the multi-dimensional matrices into chromatin states ^54,55^ using ChromHMM software (v1.26-0) with a bin size set to 200 bp. The LearnModel program of ChromHMM was used to learn the chromatin state model with the numstates being set as 3-10 and 12 states. The resulting output files for states with and without RT are available and they include bed files for genome browser visualization.

### Statistical analysis

For virtual 4C arc plots, we used plotGardner v1.0.17 ^56^ package for the R statistical suite ^57^ for the visualization of genomic regions. Two similar sized early and middle RT segments were selected as 4C baits and then their corresponding 10-kb bins were extracted. Each 10-kb bin was annotated with its RT class, and all bins in contact with the bait bins were retrieved. Hexbin plots were also drawn using the R package ggplot2 v4.1.2 ^58^ with Hi-C EV versus different chromatin marks and early-S, middle-S RT versus chromatin marks at 50-kb resolution. R scripts used for plots are available on GitHub.

### Root tip nuclei for 3D-FISH

Maize seedlings were grown and nuclei were prepared and sorted as described above for MMOA-seq. For 3D quantitative FISH with the mixed population nuclei, we combined 30 µL each of the flow-sorted EdU-labeled nuclei from flow sorting gates (Figure 1a) for G1 (1.5M/mL, 16%), early-S (1.9M/mL, 21%), middle-S (1.5M/mL, 16%), late-S (0.36M/mL, 4%), and G2 (3.9M/mL, 41%). In this mix, the S-phase nuclei were ∼41% of the total. For 3D-FISH with G1 nuclei, only flow-sorted G1 nuclei were used (Figure 7c, inset panel).

### Probe labeling for 3D-FISH

Oligonucleotide probe libraries were designed by DAICEL Arbor Biosciences using three sets of sequences as inputs, early-S, middle-S, and late-S RT segments. These annotation segments were based on the Repli-seq data from Wear et. al, (2017) ^2^, realigned to B73v5 using the Repliscan pipeline ^38^ to produce a DNA RT class annotation file, RT_class_ALL_27800_b73v5_9colBed_vhsf521e.BED. From this annotation file, we derived coordinates for two sets of genomic regions (early or middle) within the first 105 Mb on chromosome 5, which corresponds to the short arm. These were used as input to obtain two libraries of oligos that were *T*m-matched ∼45-mers, high-density, and uniformly spaced at high density across the chromosome arm. The resulting sets of oligos included a total of 22,460 or 27,390 oligos from the early-S or middle-S regions, respectively, and the coordinates of the uniquely named oligos are provided as bed files.

A series of experiments were conducted to convert these dsDNA libraries into fluorescently labeled single-stranded DNA (ssDNA) libraries. Briefly, the dsDNA library underwent PCR amplification to obtain a DNA yield sufficient for *in-vitro* transcription. The purified DNA was processed using the Qiagen QIAquick PCR Purification Kit and quantified using a spectrophotometer (NanoDrop). After DNA purification, *in vitro* transcription was carried out using the MEGAshortscript TM T7 Kit (Thermo Fisher), using 480 ng/μL DNA as template per reaction. Following transcription, RNA purification was performed using the RNA Clean-Up Mini Kit (Macherey-Nagel). RNA was quantified by NanoDrop to verify that the concentration was above 1 μg/μL, as required by the reverse transcription PCR (RT PCR) step. Single-stranded DNA (ssDNA) was generated through a reverse transcription reaction using SuperScript IV Reverse Transcriptase, with the addition of 52 μg of RNA to obtain sufficient yield of ssDNA. At this stage, ssDNA probe pools were labeled with any of three different fluorophores (Eurofins Genomics US; green, ALP_a488b: [Alexa488]; red ALP_a594a: [Alexa594]; or far red, ALP_a647n: [ATTO647N]). The process resulted in RNA-DNA hybrids and some unincorporated primers. Exonuclease-I was used to digest unincorporated primers, and RNase was used to digest the RNA from RNA-DNA hybrids. After RNA and primer digestion, labeled ssDNA was purified using the Zymo Quick-RNA purification kit and quantified.

### Fluorescent in situ hybridization

The FISH protocol was adapted from Bass *et al.* (1997) ^59^ as detailed by Howe *et al.* (2013) ^60^, using a polyacrylamide embedding technique for 3D FISH analysis. Fixed EdU-labeled nuclei were embedded in a thin layer of optically clear 3X acrylamide mix on a glass slide. Then wash buffers and prehybridization buffers were exchanged by addition and removal of 200-µL drops. A buffer containing RT class-labeled probes, 50% formamide, and 2X SSC was added and incubated at 37°C for 30 min for prehybridization. After prehybridization, incubation slides were placed on a hot plate at 65°C for 30 min for genomic DNA denaturation. After denaturation, the slides were placed back in the 37°C incubator overnight for hybridization. The following day, the slides underwent serial washing with buffers of increasing stringency. Finally, slides were mounted with mounting media and sealed with Sally Hansen Hard as Nails polish.

### Deltavision microscopy and Image analysis

Following the FISH experiment, 3D images were captured with 0.2-micron projections by multiple wavelength iterative deconvolution microscopy. The raw data was subsequently subjected to 3D iterative deconvolution and chromatic aberration correction (CAC). For analysis purposes after 3D data collection, individual nuclei with probe signals were cropped to allow quantitative signal colocalization analysis using the inbuilt solid object builder polygons method.

We used DeltaVision software (softWoRx v6.5.2) programs EditPolygon and VolumeBuilder to segment the FISH signal regions in 3D. We used EditPolygon to trace the edges of the FISH signals manually with a mouse, and VolumeBuilder to connect the polygon series into a single 3D volumetric object with a closed and continuous surface area. Using VolumeBuilder, we obtained the object volumes summed intensities (sum of all voxels within the object) for DAPI, which we used to calculate the DAPI concentration. For these analyses, the control regions were similarly built in non-FISH "bulk euchromatin" areas in the same nucleus, avoiding heterochromatic and nucleolar regions. Quantitative segmentation data analysis was performed on at least 50 nuclei for each of the early and middle probes. All 3D images were stored on the image repository omero.bio.fsu.edu, allowing for additional segmentation analysis.

## RESULTS

### Repli-seq scheme and mini-domain chromatin fiber RT model

The overall experimental scheme and replicative labeling approach that previously led to the two-compartment mini-domain RT model ^24^ is summarized in Figure 1. Maize (*Zea mays* L.) roots are labeled *in vivo* with EdU, which is subsequently clicked with Alexa Fluor 488 (AF-488), allowing for bivariate FANS sorting into three populations of S-phase nuclei, early-S, middle-S, and late-S (Figure 1a). Representative examples illustrate the canonical labeling patterns for each stage (Figure 1b), with early-S and middle-S synthesis broadly dispersed throughout the nucleoplasm, whereas late-S synthesis is limited to discrete patchy heterochromatic regions. In maize, the cytological similarity observed between the early-S and middle-S phases in their DNA labeling patterns is one of several factors that support the classification of middle-S-replicating chromatin as euchromatin ^24^.

At the molecular level, RT is visualized as Repli-seq coverage profiles from early-S, middle-S, or late-S libraries as previously described ^2^. For this study, Repli-seq data was mapped to the current maize B73 genome assembly (B73v5) and used to annotate and segment the genome into discrete DNA RT classes using Repliscan ^38^.

The maize genome comprises 10 metacentric chromosomes, each ranging from approximately 150 to 300 Mb in size. In this study, we use chromosome 5 as a representative example based on the fact that it represents a typical maize metacentric chromosome, and that its RT profiles reflect those of the whole genome as previously defined by Repli-seq ^2^.The RT profiles along the entire length of maize chromosome 5 (Figure 1c) illustrate regional enrichments for early, middle, and late replication. The early-S RT profiles show the highest coverage in the more distal regions of the chromosome arms, where gene density tends to be higher in maize chromosomes ^61^. The middle-S RT profiles show a unique pattern, being relatively evenly distributed across most of the chromosome. In contrast, late-S RT profiles show the highest coverage near the center of the chromosome at the centromeric and pericentromeric regions. A distinct non-centromeric block of late-S replicating chromatin is also seen at the heterochromatic knob, embedded in the long arm of chromosome 5. To gain further insight into the relationships among the different RT classes, we inspected 2-Mb non-heterochromatin regions on chromosome 5 to specifically focus on the distribution of early-S versus middle-S (Figure 1d, blue and green) RT classes. Zooming in on 2-Mb regions outside of the centromeric heterochromatin illustrates a striking pattern of RT, in which closely interspersed alternating early-S and middle-S segments are evident, switching back and forth every ∼100 kb. This mini-domain pattern is notably distinct from the large Mb-size RT domains commonly described for mammals ^62,63^.

A genomic composition analysis of early-S, middle-S, and late-S RT regions revealed that all three classes were predominantly composed of transposable elements (TEs), which comprise ∼85% of the maize B73 genome ^61^. Given their abundance and distribution across all three RT classes (Figure 1e), it is clear that TEs or repeats alone are not a defining feature of RT-associated genome organization. We wanted to further explore the spatial organization of the genome with respect to RT regions, specifically focusing on early-S versus middle-S. The previously proposed "mini-domain chromatin fiber RT model" ^24^ is illustrated (Figure 1f-h).

According to this model, maize euchromatin, which is not uniformly stained in cytological preparations (DAPI image, Figure 1f,g), exists in two closely interspersed subcompartments that are evident in S phase, corresponding to early-S and middle-S replicating regions. The early-S chromatin regions are relatively weakly stained with DAPI, whereas the middle-S chromatin regions exhibit stronger DAPI staining, as depicted in the interpretive diagram (Figure 1h). We aimed to test whether the mini-domain model, originally defined based on two distinct chromatin types during S phase in root tip cells, represents a broader principle of maize nuclear architecture. Specifically, we investigated whether this model applies not only to nuclei in S phase but also throughout interphase and across different maize tissues. This idea was first tested using high-throughput conformation capture (Hi-C) analysis with nuclei from the entire mitotic cell cycle (Supplementary Figure 1a-c).

### Hi-C data reveals covariance and correlation with early and middle RT in maize euchromatin

We used formaldehyde-fixed root tip nuclei from 3-day-old seedlings to prepare Hi-C libraries in biological replicates of 1.25 million nuclei each (Supplementary Figure 1a-d). For comparison, we also made Hi-C libraries from immature earshoot nuclei. The analysis of the resulting Hi-C libraries provided genome coverage and valid pair contact frequencies comparable to that previously reported for maize bundle sheath cells ^41^. This confirmed that the contact patterns remained intact following the nuclei isolation, flow sorting, and library preparation steps (Supplementary Table1).

To compare nuclear architecture to DNA RT, we performed nuclear compartment analyses using Hi-C as shown in Figure 2. To examine early-S versus middle-S, we looked at three different non-heterochromatin 2-Mb regions from the short arm of chromosome 5. We plotted the RT profiles and segments along with the chromosome-wide eigenvector (EV) and insulation score (IS) analyses with a 50-kb bin size for all four biological replicates (Figure 2b-d). The IS of a genomic region measures its frequency of interaction with the neighboring regions. A low or negative IS indicates a scarcity of local interactions relative to longer-range contacts ^49^.

**Fig. 2.**
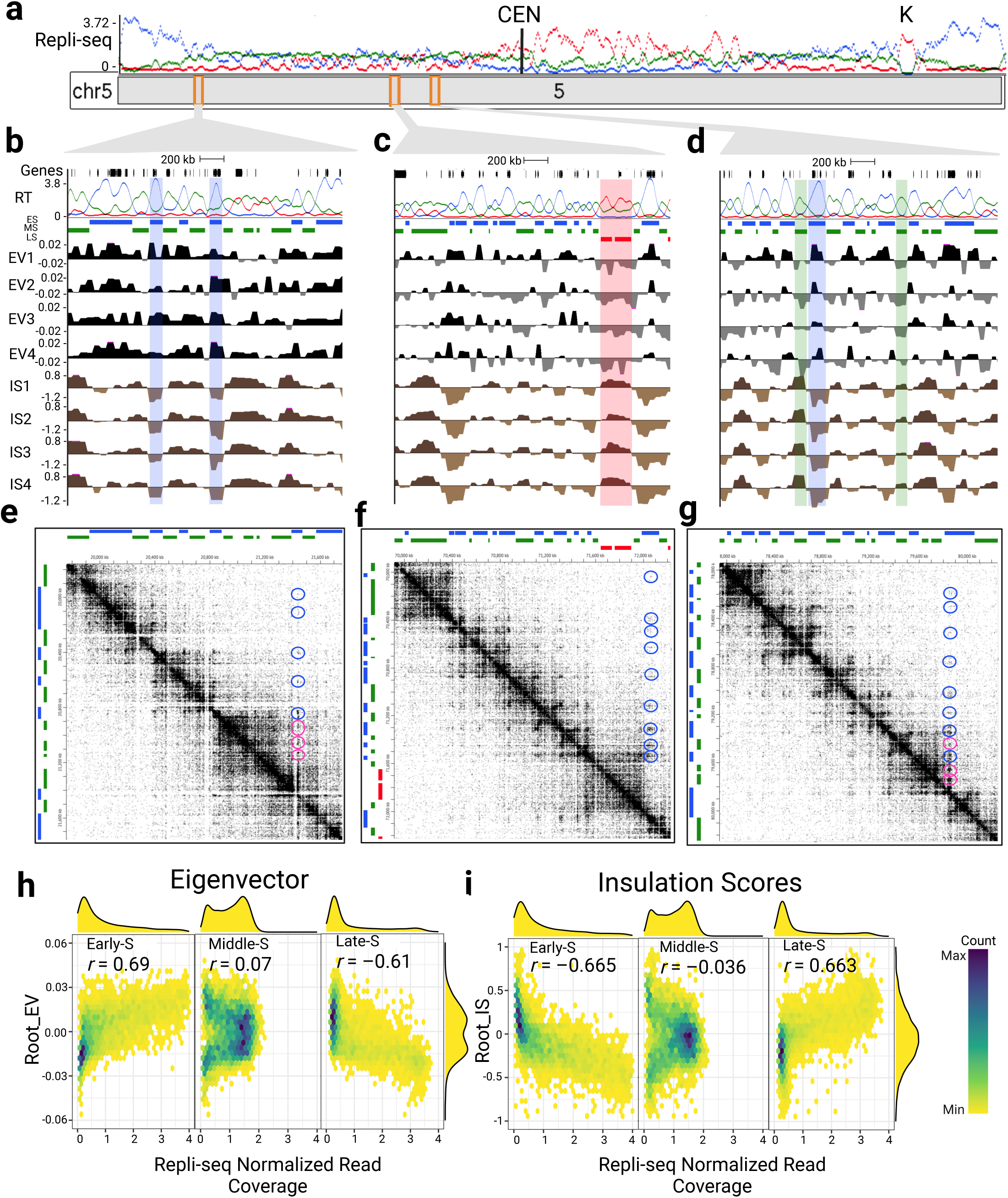
Hi-C on maize root tip nuclei. **(a)** Repli-seq profile across chromosome 5, with the knob (K) and centromere (CEN) labeled. **(b-d)** UCSC Genomaize browser views of three 2-Mb regions (indicated by orange rectangles), showing the genes, RT profiles, chromosome-wide eigenvector (EV) analysis, and insulation score (IS) tracks for four biological replicates. Light blue, green, and red shaded regions correspond to early, middle, and late RT segments, with their respective EV and IS patterns. **(e-g)** Hi-C contact matrices of the 2-Mb genomic regions in panels b,c, and d, respectively. RT classification segments (early;blue, middle;green and late;red) are shown on the top and left edges of the matrices. The color intensity represents the frequency of contacts between two loci. Blue and pink circles highlight examples of early-early and early-middle contacts, respectively. **(h)** Correlation plots of root tip Hi-C eigenvector (y-axis) and root tip Repli-seq early-S, middle-S and late-S data (x-axis). **(i)** Correlation plots of root tip insulation score data (y-axis) and root tip Repli-seq data (x-axis). Hexagonal cell intensity indicates the number of data points (scale on the right), and density plots on the top and left show independent distributions for each variable.

Strikingly, most of the points at which the EV and IS values switched from positive to negative mirrored the locations where RT segment switches occurred (Figure 2b-d). For instance, the light-blue shaded regions, representing early-replicating segments, showed positive EV and negative IS values. The low IS of early-S RT regions denote a greater tendency for long-range contacts. Relatedly, the green-shaded regions, indicating middle-S RT segments, correspond to negative EV and positive IS values, suggesting more confined interactions ^49^. Finally, the red-shaded areas show late-S RT regions with corresponding negative EV and positive IS values.

These Hi-C patterns uncover smaller-scale nuclear architectural features that extend beyond the global A (early) and B (late) compartments by differentiating early-S and mid-S euchromatin regions. We next pooled the root tip biological replicates based on their similarity (Supplementary Figure 2a) and used the pooled data to generate the contact matrix plots that are shown in relation to the Repliscan segment classification (Figure 2e-g). The contact matrices showed a strong diagonal signal for the expected abundant interactions among adjacent loci. Beyond the diagonal, interaction hotspots were observed (examples highlighted in blue and pink circles). Blue circles highlight contacts between two non-adjacent early RT regions, while pink circles mark interactions between early and middle RT regions. These distal contact hotspots represent long-range interactions within these representative 2-Mb regions.

Notably, early-S replicating segments appeared to display more long-reaching interactions with other early-S replicating segments compared to adjacent middle-S replicating segments. Some of these early-to-early contacts extend beyond 1 Mb, skipping multiple intervening RT segments consistent with the low IS of early-S RT regions (Figure 2b-d).

To determine if these relationships between nuclear architecture and RT hold true for the rest of the genome, we carried out a correlation analysis between RT and Hi-C. For this we combined the chromosome-wide Hi-C data (EV and IS) from all ten chromosomes. The early-S Repli-seq coverage values showed a strong positive correlation (*r* = +0.69) with the eigenvectors, whereas the middle-S Repli-seq coverage showed a weak positive correlation (*r* = +0.07). The late-S Repli-seq coverage values showed a strong negative correlation (*r* = -0.61) with the eigenvectors (Figure 2h). When doing a similar genome-wide correlation with IS, we observed a strong negative correlation with early-S Repli-seq coverage (*r* = -0.67), a weak negative correlation with middle-S coverage (*r* = -0.04), and a strong positive correlation with late-S Repli-seq coverage (*r* = +0.66) (Figure 2i). For both EV and IS, the strongest but opposite correlations were observed for early-S compared to late-S chromatin. The middle-S chromatin showed the same direction of correlation as that of the early-S chromatin, but with much smaller correlation values. Together, these data show that genomic regions distinguished by their timing of replication also correlate with distinct Hi-C-defined chromatin and genomic organization across the whole genome.

To examine the impact of resolution on the observed correlation patterns, we performed similar correlation analyses at multiple resolutions. Chromosome-wide eigenvector (CEG) analyses at 25 kb and 100 kb, as well as block-based eigenvector (BEG) analysis at 50 kb, confirmed similar (although slightly weaker for BEG) trends across all resolutions at both the chromosome and genome-wide levels (Supplementary Figure 3). In addition to EV analysis, we conducted IS analysis at a range of resolutions (1 kb, 10 kb and 25 kb), using two sliding window sizes (2.5 Mb and 1 Mb; Supplementary Figure 4). All of these additional IS analyses have similar correlation with our RT data as observed at 50 kb CEG analysis.

We next compared our Hi-C data for root tips to the EV and IS data from earshoots analyzed in parallel and to maize bundle sheath cells from a published study ^41^. Pairwise comparisons between tissues (Supplementary Figure 5) revealed strong genome-wide similarity between root tip and earshoot Hi-C profiles (*r* = 0.85 for both EV and IS), both of which are actively growing tissues. We also observed slightly weaker but significant correlations between root tip and bundle sheath Hi-C profiles (*r* = 0.73 for EV and 0.66 for IS), possibly reflecting some level of cell-type independent global conservation of genome organization.

We also investigated whether any of the ten maize chromosomes showed more or less tissue-specific variation compared to the whole genome (Supplementary Figure 6). Interestingly, we found that the root tip and earshoot Hi-C data showed a strong positive correlation across all chromosomes with the notable exception of chromosome 6, which contains the nucleolar organizing region near the end of the short arm. The correlation coefficient for chromosome 6 was *r* = 0.62, compared to all the others which ranged from *r* = +0.78 to +0.95 (Supplementary Figure 6a). We found a similar situation for root tip versus bundle sheath, with chromosome 6 showing the lowest correlation (Supplementary Figure 6b). In addition, most of the other chromosomes showed relatively weaker correlations with bundle sheath data, likely due to variations in Hi-C library preparation or cell type or both. Using a stratum-adjusted correlation coefficient (SCC) analysis ^64^, we also found that root tip to earshoot similarity was higher than that of either one compared to bundle sheath (Figure 3a).

**Fig. 3.**
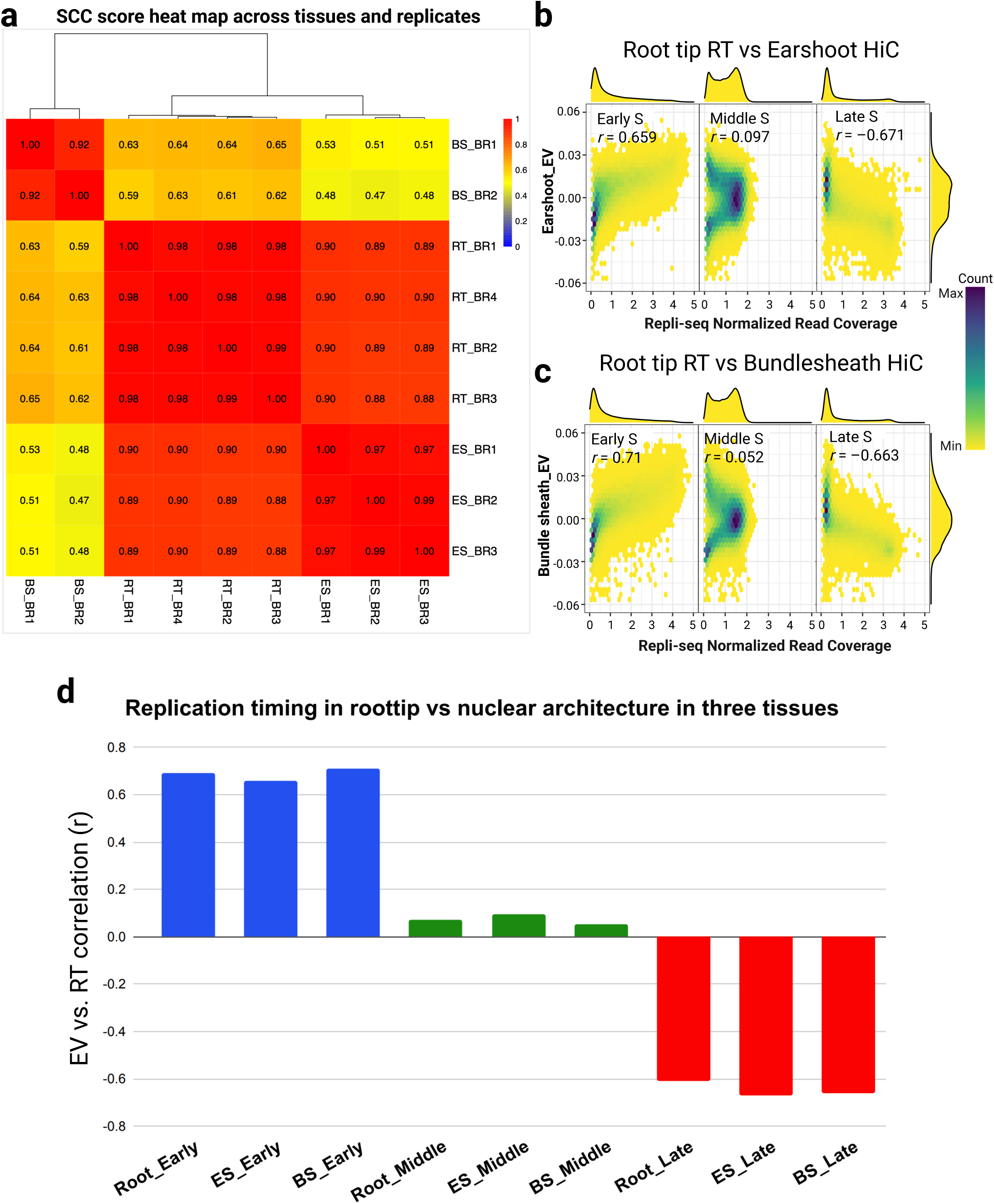
Root tip Repli-seq versus Hi-C from different tissues. **(a)** A stratum-adjusted correlation coefficient (SCC) score heatmap across biological replicates of the same and different tissues showed the reliability of multiple replicates. **(b)** Correlation plot showing root tip Repli-seq data (x-axis) and earshoot eigenvector (EV) analysis data (y-axis). **(c)** Correlation plot showing root tip Repli-seq data (x-axis) and bundle sheath eigenvector (EV) analysis data (y-axis). Density plots above and to the left of each plot represent the independent distributions of the variables. **(d)** A bar chart summarizing the correlation analyses comparing Repli-seq data (early;blue, middle;green, late;red) in root tip versus Hi-C eigenvector data in root tip, earshoot, and bundle sheath tissues.

Given the strong correlations between RT and Hi-C defined nuclear architecture in root tip, we wondered if the RT classification alone from root tip could be predictive of Hi-C nuclear architecture in other tissues. For this, we did correlation analysis between root tip Repli-seq and the Hi-C EV data of the other two tissues, for which we do not have RT data. These results are summarized in Figure 3. The root tip Repli-seq-defined genome structure closely mirrored the Hi-C eigenvectors of the other tissues (Figure 3b-c), which can be seen locally (Supplementary Figure 7b-c) and globally (Figure 3d). These analyses demonstrated that RT in root tip predicts chromatin structure across other tissues, including earshoot and bundle sheath, suggesting a robust association between RT and chromatin organization across the plant.

### Hi-C contacts are enriched within RT classes

We used the pooled Hi-C valid contacts to examine the degree to which moderate to long range contacts were occurring between regions with the same RT assignments, i.e. within-class contacts, as predicted by the two-compartment mini-domain model for euchromatin (Figure 4). We assigned RT class annotations to each of the two members of the valid pair contacts and calculated the frequencies of contacts with matching and differing pairs for early-S, middle-S, or late-S regions. The analysis revealed that of all the valid pair contacts, early-early accounted for 28% (2.93 million), middle-middle for 24% (2.47 million), and late-late for 16% (1.65 million) (Figure 4a). Contacts between two different RT classes were less frequent, with 16% early-middle, 13% middle-late, and only 3% early-late. These findings indicate the "same-class" categories are the most frequent, even after normalizing to account for their relative abundance (Figure 4b).

**Fig. 4.**
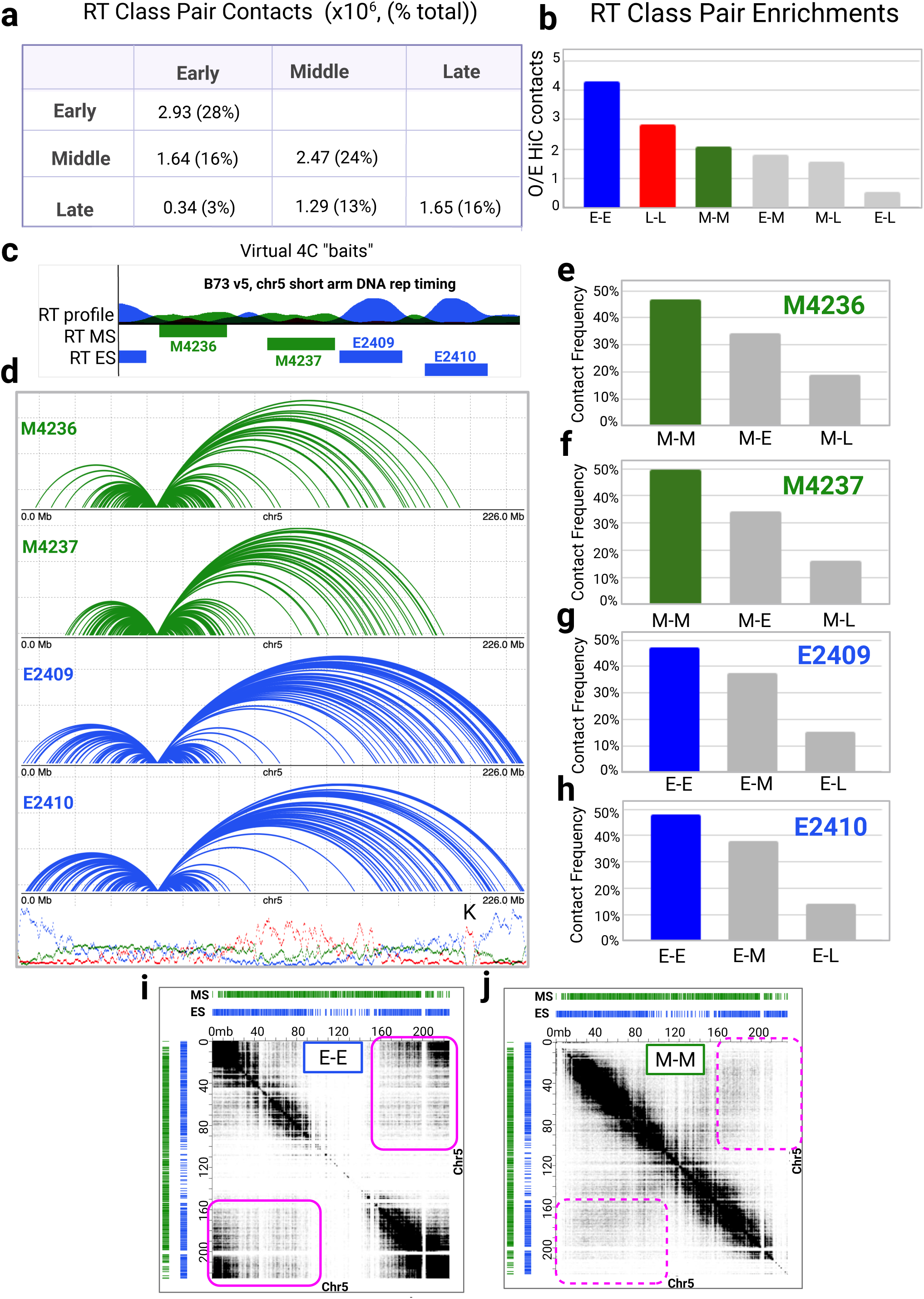
RT class contact enrichment and virtual 4C analysis. **(a)** Table showing the contacts for each RT class pair in number per million and percentages of valid Hi-C contact pairs. Contacts between different RT classes in both directions (e.g., early-middle or middle-early) are represented within the same cell. **(b)** Observed versus expected Hi-C valid contacts, with the same RT class contacts shown in blue (early-early), green (middle-middle), and red (late-late). Gray bars represent different RT class contacts. **(c)** UCSC Genomaize browser view of virtual 4C baits targeting early and middle RT segments, below the RT profile from the short arm of chromosome 5 region chr5:60,579,859-61,470,739. **(d)** Contact maps extending across chromosome 5 of each of the four bait regions shown in panel c. Blue and green arcs represent genomic contacts originating from each early and middle RT segment, respectively. The height of lines are autoscaled on the Y axis according to the longest contact in the plot. The RT profile for the whole chromosome is shown at the bottom of the plot. **(e-h)** Bar charts displaying the contact pair frequencies per bait segment. **(i)** Hi-C contact matrix showing only early-early RT contacts for all of chromosome 5. **(j)** Hi-C contact matrix showing only middle-middle RT contacts for chromosome 5. The contact matrix plots are displayed with settings of 1-mb resolution and 40 color intensity. Contrasting regions illustrate long-distance contact frequency differences between M-M (dashed pink boxes) and E-E (solid pink boxes) in the contact matrices.

To examine more closely the same-RT-class contacts, we conducted a virtual 4C analysis of two early and two middle RT genomic regions, each approximately 150 kb, located on chromosome 5 (Figure 4c). This approach allowed for the identification of long-range interactions centered around a specific genomic region of interest, known as the “bait” region, or virtual bait in this case. Two early and two middle RT segments were designated as baits, and their same-class contacts were plotted along the entire chromosome 5 (Figure 4d). All four baits showed long distance contacts, including to regions on the other arm of the chromosome, but the early-early contacts extended further, even beyond the distal knob on the long arm of chromosome 5. This difference can not be explained by the location of the baits, because they come from the same local region (Figure 4c), which represents less than 1% of the 226 Mb chromosome 5. The per-bait analysis also showed that the same class contacts were the predominant type. The least frequent contacts were between early or middle baits and late-S regions for all four segments analyzed (Figure 4e-h). We observed consistent results when we decomposed the contact matrices for all of chromosome 5 into early-to-early (Figure 4i) and middle-to-middle RT contacts only (Figure 4j).

We next investigated the persistence of these same-class contact tendencies as a function of distance relative to the linear genome. The results demonstrated the expected distance-dependent decay of all contact classes, with early-early RT class contacts showing the highest frequency at any given distance (Supplementary Figure 8a). This enrichment was not a result of the relative abundance of the RT classes per se, as shown using a control with the RT positions randomly shuffled (Supplementary Figure 8b).

Overall, the contact class pair enrichments and the virtual 4C bait results showed that same-class contacts were more frequent than cross-class contacts, and that early RT segments exhibit more of the longest-range interactions than the middle RT segments.

### Hi-C-based chromatin architecture and Repli-seq aligns with epigenomic and transcriptional features

To test how our Hi-C data correlated to genomic and genetic features, we checked the correlation of our 50-kb Hi-C eigenvectors with epigenetic and transcriptional features known to be associated with chromatin structure (Figure 5). We first examined how the Hi-C data correlated with histone post-translational modifications using ChIP-seq data from B73 maize root tips ^2,65,66^.

**Fig. 5.**
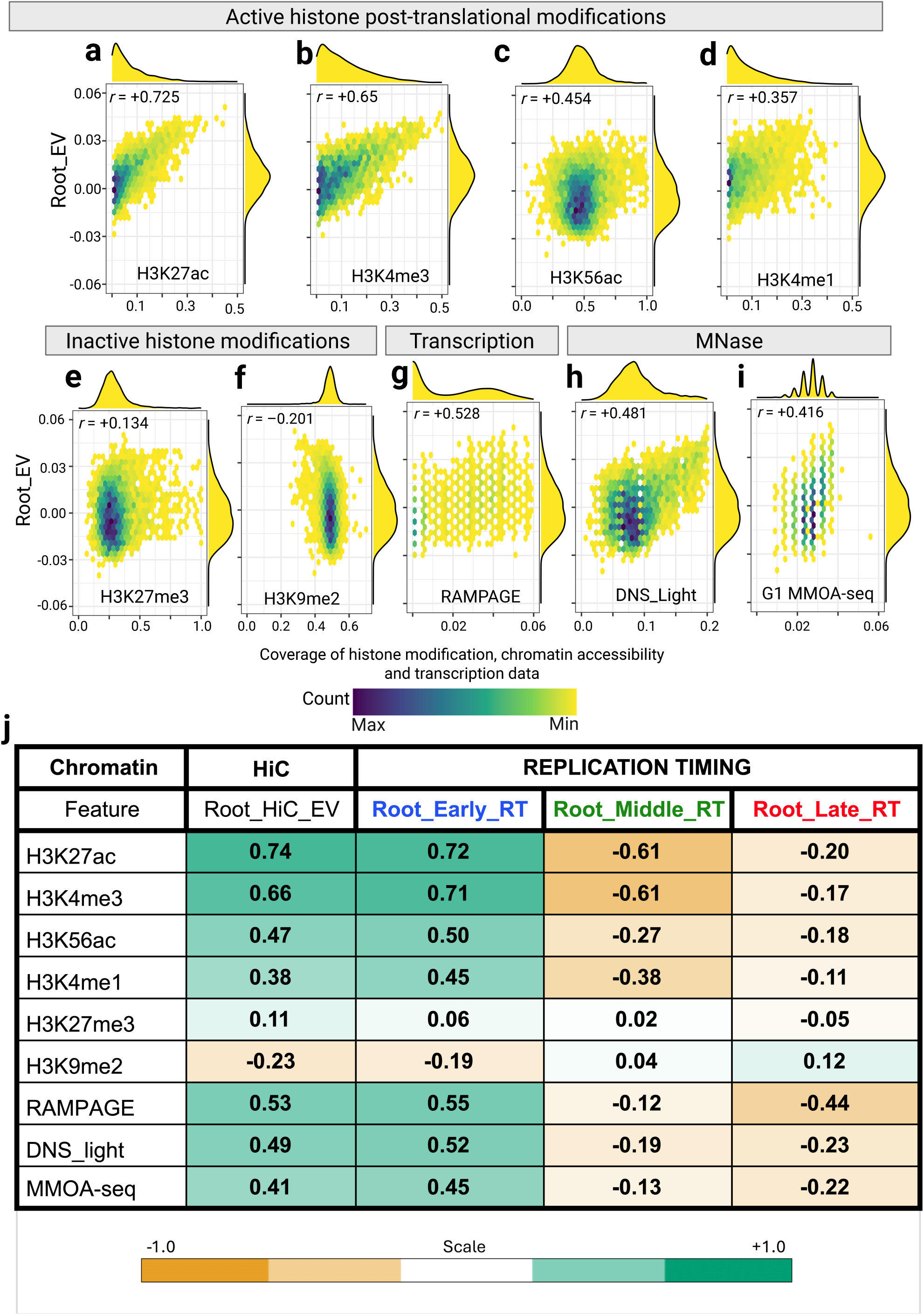
Hi-C correlation with other chromatin features. (a-f) Correlation plots between root tip Hi-C data and histone post-translational modifications: **(a)** H3K27ac, **(b)** H3K4me3, **(c)** H3K56ac, **(d)** H3K4me1, **(e)** H3K27me3, and **(f)** H3K9me2. **(g)** Correlation plot between root tip Hi-C data and RAMPAGE transcriptional data. **(h-i)** Correlation plots between root tip Hi-C data and chromatin accessibility: **(h)** root tip differential nuclease sensitivity (DNS) light digest data and **(i)** G1 MMOA light digestion. **(j)** Correlation values between different chromatin features and Hi-C data, early RT, middle RT and late RT Repli-seq data. The H3K27ac, H3K4me1, H3K4me3 and RAMPAGE data was taken from Cahn *et al.* (2024) ^66^, H3K56ac and H3K27me3 data from Wear *et al.* (2017) ^2^, H3K9me2 data from Wear *et al.* (2020) ^65^, DNS light digest data from Turpin *et al.* (2018) ^53^ and MMOA-seq data from this paper.

We previously anticipated that modifications associated with active genes would be positively correlated with early-S RT regions within euchromatin, whereas repressive marks would not be or less so ^27^. Indeed, we found significant correlations between our root tip Hi-C eigenvector (EV) data analyzed at 50-kb resolution on chromosome 5 and the active gene-associated marks (Figure 5a-d). Among the marks with positive correlations, H3K27ac and H3K4me3 had the strongest correlations with *r* = +0.73 and +0.65, respectively. The values for the histone marks and the eigenvector varied along the length of the chromosome, showing a general trend of higher values in the chromosome arms for all of the marks tested, except for H3K9me2 (Supplementary Figure 9a-d). Consistent with our findings, regions enriched for H3K56ac (EV *r* = +0.45) and H3K4me3 (EV *r* = +0.65) were previously classified as early-S replicating regions, which also show enrichment for genes with high median expression levels ^2^. Regions enriched for H3K27me3, on the other hand, exhibited a weakly positive association with Hi-C EVs (Figure 5e, *r* = +0.13). The H3K27me3 mark is known to be associated with poised genes ^67,68^ and also found to be enriched in middle-S replicating regions and genes with lower median expression levels ^2^. The only histone mark we examined that showed a negative correlation with our root tip EV was H3K9me2 (Figure 5f, r = -0.20), which is known to be associated with constitutive heterochromatin ^69,70^. H3K9me2 is typically associated with transposable elements (TEs) which are broadly distributed across the whole maize genome consistent with the distribution observed across chromosome 5 (Supplementary Figure 9f).

We also investigated the correlation between Hi-C EVs and transcriptional activity estimates from the RAMPAGE transcription start site mapping data and it showed a positive correlation (Figure 5g, *r* = +0.53) ^66^. The correlation plots for the RAMPAGE data show substantial dispersion, which we attribute to the natural sparsity of the data, which is at single base resolution. Finally, we compared Hi-C EV with chromatin accessibility using micrococcal nuclease (MNase) sensitivity profiling datasets from maize root tip nuclei. The read coverage levels from a light MNase digestion ^53^ showed a positive correlation of *r* = +0.48 with Hi-C EVs (Figure 5h, Supplementary Figure 9h). Likewise, the open chromatin defined by sequencing small fragments from light MNase digests in purified G1 nuclei showed a positive correlation of *r* = +0.42 with Hi-C EVs (Figure 5i, Supplementary Figure 9i). These were calculated for chromosome 5 because it was selected for additional cytological analysis, but similar trends were observed genome wide for correlations between these same epigenomic features and both Hi-C and RT (Figure 5j).

Given that RT is intimately coupled to a multitude of chromatin features, we tested the effect of including Repli-seq data inputs for chromatin state (CS) analyses. For this analysis, we performed high-resolution (200 bp) chromatin states analysis using ChromHMM (Ernst & Kellis, 2010), employing two input sets: one combining eight chromatin features with three Repli-seq datasets, and the other with the same eight features excluding Repli-seq data. The eight common chromatin features used in both training sets included ChIP-seq for H3K27ac, H3K4me3, H3K4me1, H3K56ac, H3K27me3, and H3K9me2, plus RAMPAGE for transcription and light digest MNase for open chromatin. Among the tested models, from 3-state to 12-state, we selected the 6-state models for further analysis based on the biological interpretability and the transition parameters, which provide a measure of state resolution quality (Supplementary Figure 10). The inclusion of Repli-seq resulted in a more balanced distribution of chromatin state abundances (Figure 10a,b). Including Repli-seq data as input, CS1 and CS2 are distinguished by the absence (CS1) or presence (CS2) of active gene histone marks, H3K4me3, H3K4me1, and H3K27ac. The middle-S replicating regions are similarly distinguished by the absence (CS4) or presence (CS3) of these same active gene histone marks (Supplementary Figure 10c-e).

We were able to integrate RT data at high resolution along with epigenomic features of chromatin structure, using transition state data. We found that including Repli-seq data improved the resolution across the entire range of trained models (Supplementary Figure 10i). Overall, we conclude that the ChromHMM better resolves chromatin states when RT data is included. Similar to the replicative labeling patterns (Figure 1b), the tendency for early and middle S in similar chromatin states (see emission heatmaps of 3, 4 and 5 state models; Supplementary Figure 12) provides additional evidence that these two RT classes represent functionally unique sub-types of euchromatin (Supplementary Figure 12, and compare Supplementary Figure 10c CS2 to CS3).

### 3D cytological evidence for the spatial bifurcation of euchromatin into early-S and middle-S chromatin regions

Having shown that Hi-C data aligns well with the two-compartment mini-domain model (Figure 1h), we next explored a different type of evidence, 3D cytology, using a chromosome painting strategy based on probes designed to detect early-S and middle-S RT sequences within fixed nuclei (Figure 6). We found that these two RT class frequencies for the short arm of chromosome 5 (chr5S) closely matched those of the whole genome (Supplementary Figure 13a). For instance, early-S segments were 30% of chr5S and 30% of the whole genome, whereas middle-S segments were 38% and 35% of chr5S and the genome, respectively.

**Fig. 6.**
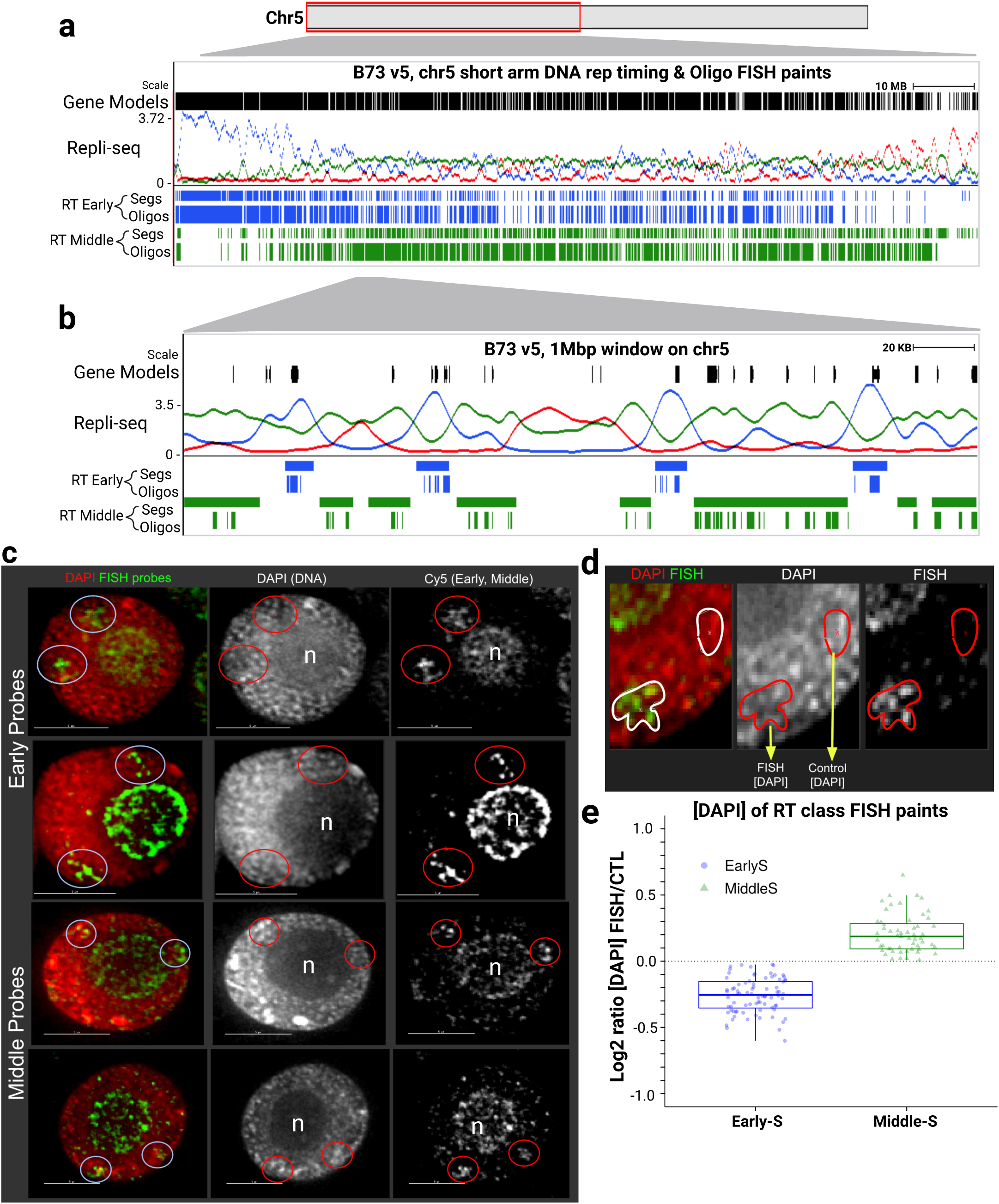
Probe design and 3D cytology of single labeled FISH. (a, **b)** UCSC Genomaize browser views of chromosome 5S **(a)** and a 1-Mb zoomed-in region **(b)**, showing gene models, Repli-seq profiles, RT segments for early (blue) and middle (green) and their corresponding 3D FISH probe oligo pools. **(c)** Cytology of total nuclei from root tips hybridized with early-S and middle-S FISH probes labeled with far-red fluorophores. Images were collected using 3D deconvolution microscopy, corrected for wavelength-dependent chromatic aberration, and displayed as separate grayscale (DAPI or FISH probes) or as merged color overlays (DAPI in red, FISH probes in green). Each row represents a single nucleus, showing intensity-averaged projections spanning 1 µm (5 Z-sections) that include the FISH signals. Total DNA was visualized in the DAPI channel, and FISH probe signals were captured as AF-647 fluorescence in the Cy5 channel. Two representative examples are shown for early-S probes (1st two rows) and middle-S probes (last two rows). The location of nucleoli is marked as "n". Regions where FISH probe signals occur are indicated by circles in grayscale images (red circles) or color overlays (blue circles). The two similarly sized signal clouds represent the chromosome 5 homologs. All scale bars = 5 µm. **(d)** Individual nuclei with FISH probe signals were cropped for quantitative colocalization analysis using the inbuilt solid-object builder polygons method. The DAPI density of FISH signal regions was compared with non-FISH euchromatin control regions from the same nucleus. **(e)** Box plot showing the log2 ratio of DAPI density in early probe/control (∼80 probe signals) and middle probe/control (∼55 probe signals) regions. This box plot contains data from a mixed population of G1, S and G2 nuclei. The ratios were statistically different according to two-sample unequal variance (Welch’s t-test, p=1.34E-28).

**Fig. 7.**
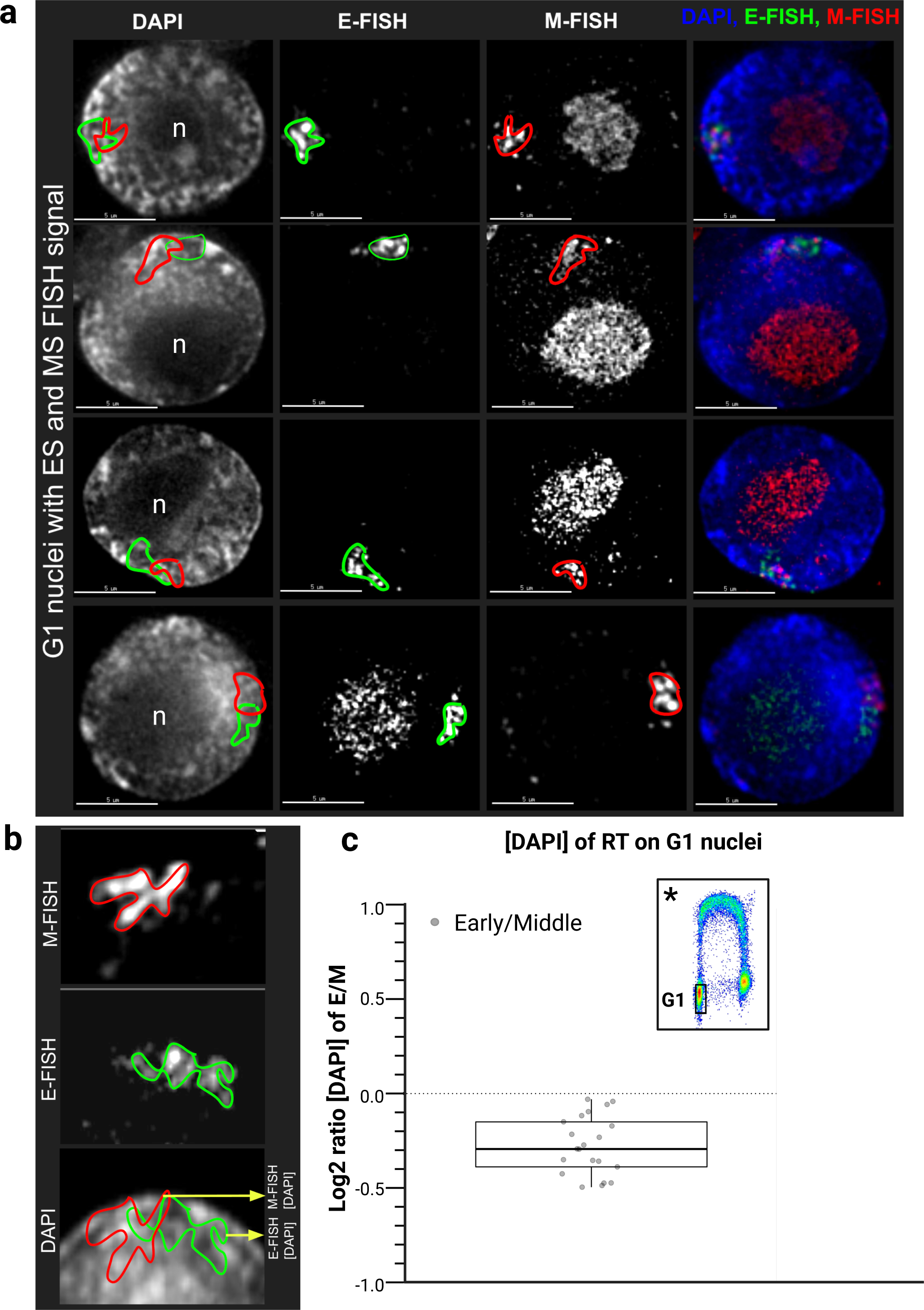
3D cytology of two color FISH. **(a)** Cytology of G1 root tip nuclei with total DNA stained with DAPI and hybridized with early-S and middle-S FISH probes labeled with green (AF-488, FITC channel) and far-red (AF-647, Cy5 channel) fluorophores, respectively. Images were acquired using 3D deconvolution microscopy, corrected for wavelength-dependent chromatic aberration, and displayed as grayscale (DAPI, FITC, or Cy5 channel) or as merged color overlays (DAPI, blue, early-FISH in green, middle-FISH in red). Each row represents a single nucleus, showing intensity-averaged projections spanning 1 µm (5 Z-sections) that include the FISH signals. Four representative examples of dual-labeled G1 nuclei are shown. The location of nucleoli is marked as "n". Green and red outlines on grayscale images highlight early-FISH (E-FISH) and middle-FISH (M-FISH) signals and the same regions on the DAPI image where these signals are present. **(b)** Individual nuclei with FISH probe signals were cropped for quantitative colocalization analysis using the inbuilt solid-object builder polygons method. The DAPI concentration of early-FISH and middle-FISH probe signal regions were directly compared. **(c)** Box plot displaying the log2 ratio of DAPI concentration in early-FISH/middle-FISH regions in the same nucleus. Bivariate plot inset (*) in the corner highlights the G1-gated nuclei population used for this FISH experiment (for plot details see Supplementary Figure 14).

Therefore, we designed RT-class-specific oligo FISH painting probes at comparable densities for early-S and middle-S spanning the entire chr5S (Supplementary Figure 13b). These collectively account for 68% of chr5S, and the probe distributions match those of the RT classes (Figure 6a). These are better seen by focusing on a 2-MB region from the middle of chr5S (Figure 6b).

For this RT class oligo FISH painting experiment, early-S and middle-S probes were labeled with different fluorophores and hybridized to a mixed population of nuclei from the entire cell cycle, made by pooling G1, early-S, middle-S, late-S and G2 nuclei (Supplementary Figure 14d). Representative images from 3D deconvolution image datasets are shown as 1-µm Z-stack projections, a five-optical-section projection spanning the FISH probe signal regions. For quantitative analysis, we selected nuclei in which two similar sized paint signals were observed for diploid chromosome 5 (Figure 6c, circles). In these nuclei, non-specific background staining of the nucleolus (labeled "n") was commonly seen, but the maize nucleolus organizing region (NOR) is not located on chromosome 5 and this non-specific signal was disregarded. To quantify the DAPI signal in the FISH-painted regions, we manually segmented the FISH paint signals to create volumetric polygon objects to obtain DAPI density (average photon counts per cubic micron). As a control, random non-FISH euchromatin regions were selected, avoiding the nucleolus and bright knobs (described in Methods). In this way, we could obtain the DAPI values for chromatin in painted RT class probe regions relative to bulk euchromatin control regions (Figure 6d). By plotting the log2 ratio of DAPI density in the FISH painted versus control regions, we showed that the average DAPI concentration in early-S chromatin (mean of -0.26) was less than the control regions, whereas the middle-S FISH chromatin (mean of +0.21) was higher. The ratios of RT painted versus control region values do not overlap for early-S versus middle-S probe sets (Figure 6e, box plot). Given that observation, along with the fact that we sampled nuclei from a mixture of G1, S, and G2 in which S phase was only ∼41% of the total, we concluded that this relationship between RT and DAPI density holds true for nuclei across the cell cycle, not just in S phase (Figure 6e).

To more definitively establish the spatial bifurcation of euchromatin outside of S phase, we performed dual probe FISH experiments using early-S and middle-S probes on purified G1 nuclei as summarized in Figure 7. Representative images of dual-labeled G1 nuclei (Figure 7a) showed spatial separation of early-S and middle-S probe regions corresponding to the same unreplicated chromosome. We repeatedly observed paired but spatially separated signals for the early-S versus middle-S probes, in contrast to their intermixed, linear arrangement along the chromosome (Figure 1d). By segmenting the FISH signal regions, we could directly compare DAPI concentration in early-S chromatin (green outlines) relative to middle-S (red outlines) chromatin from the same chromosome, avoiding any uncertainty in selecting "control" regions from other parts of the nucleus. We quantified the DAPI intensity for each chromosome 5S-based pair of FISH signals (Figure 7b), providing a log2 ratio that captures the relative chromatin compaction of the two FISH-delineated areas. We found that the average DAPI concentration ratios for all the early/middle FISH region pairs was 0.86 (sd 0.07) in the G1 nuclei (Figure 7c; expressed as log2 ratio), clearly establishing that RT defined regions are spatially separated and differentially condensed, even outside of S phase.

These results validate the mini-domain chromatin fiber RT model, demonstrating that maize euchromatin exists as an interspersed mixture of two compartments distinguishable by condensation state and RT. Early-S FISH probes localize preferentially in regions with low DAPI signals, whereas middle-S FISH probes coincide with regions of higher DAPI intensity. 3D dual-label RT FISH painting in G1 nuclei provided compelling evidence, consistent with Hi-C data, that the two euchromatin compartments are integral features of nuclear organization.

## DISCUSSION

This study sheds light on the close relationship between DNA replication timing (RT) and chromatin organization in the maize genome, focusing particularly on nuclei in the terminal 1 mm of actively growing root tips and their euchromatin compartments. By using two orthogonal methods for defining nuclear architecture, Hi-C, and 3D FISH, we demonstrated how RT reveals a spatial bifurcation of maize euchromatin (the global A) into separate early-S and middle-S compartments. These findings support our earlier mini-domain chromatin fiber RT model ^24,27^ and extend it into G1 (Figure 7a-c), outside of S phase.

Other studies in eukaryotic systems point to the fact that simple global A and B compartments are widely conserved, but are insufficient to understand the complexity of structural and functional organization of chromatin. Mammalian chromatin, for instance, has been divided into a variable number of states or subcompartments, ranging from three to eight or more, based on various approaches including epigenomic marks, modularity-based analyses, trans contacts, hierarchical clustering of domains, and data from TSA-seq, DamID, and Hi-C ^12,71–74^. Similarly, plant chromatin, primarily from Arabidopsis studies, has also been shown to possess multiple chromatin states defined by distinct features ^21,75–77^. These studies tend to agree that the global A compartment can be split into two subcompartments, A1 and A2, which replicate in that order in plants and animals ^12,41,73^. We speculate that the maize early-S and middle-S chromatin described here reflect an A1 and A2 type of subdivision, which a single assay, RT profiling, can be used to map. Consistent with this, a recent light and EM cytological analysis of the very large plant genome, *Nigella damascena,* also revealed that early-S versus middle-S replication partitioned into more open versus less open chromatin, respectively ^23^.

Here, we show fine scale interspersion of replicon-size chromosome segments with strikingly different RT, and that those timings correlate well with both Hi-C and 3D molecular cytology (Figures 2, 6, and 7). This type of finely interspersed early-S versus middle-S chromatin exhibits a clear tendency to alternate correspondingly with Hi-C measures of chromatin structure. The close interspersion of these patterns highlights differences in RT and chromatin features within what has previously been regarded as undifferentiated euchromatin.

Evidence for the nonrandom spatial distribution of early, middle, and late RT domains is supported by genomic RT profiling with Repli-seq ^2^ and 3D quantitative microscopy. Within S phase, using replicative labeling, we know that early replication predominantly occurs in distal, gene-rich euchromatic regions with open chromatin features, whereas middle replication corresponds to denser euchromatin associated with repetitive elements and less active genes^24^. We now show that the close interspersion of early and middle RT domains (e.g., Figure 1b) correlates with alternating chromatin states and persists in interphase nuclei.

We have also documented the robust covariance between chromatin interactions and RT annotations. Early RT regions exhibit higher contact frequencies with other early regions, even across long genomic distances, as evidenced by the eigenvector (EV) and insulation score (IS) analyses (Figures 2 & 4). Middle RT regions exhibit a lesser tendency for long reaching interaction (Figure 2e-g; Figure 4d, j) and show a more condensed chromatin structure compared to early RT regions (see Figure 6e, Figure 7c). These "same-RT-class" interactions may reflect different functions. For example, early-to-early interactions might be largely based on transcriptional activity, while middle-to-middle interactions, occurring in regions with fewer active genes and more abundant retroelements might reflect a chromatin packaging function.

In general, it is known that the chromatin regions that replicate at different time points in S phase, such as early versus late, also show distinct enrichments for epigenomic features in plants and animals ^2,18,78^. For instance, early-S replicating regions generally exhibit open and accessible chromatin, enrichment for active chromatin marks such as histone acetylation, and relatively higher levels of transcriptional activity. On the other hand, late-S replicating regions generally exhibit the opposing features such as closed and repressive chromatin marks, including histone methylation on H3K27 or H3K9, and DNA methylation.

In contrast to the established binary framework of S phase, which divides RT into early and late stages, our study highlights a tripartite RT model consisting of early-, middle-, and late-S phase. We focused specifically on the early and middle stages, as both exhibit broad nucleoplasmic distribution and map to the global Hi-C-defined A compartment, yet they display distinct molecular features. Notably, the chromatin state CS3 (Supplementary Figure 10c) includes middle-S and active epigenomic marks, similar to the chromatin state CS2, which includes early-S and active epigenomic marks (Supplementary Figure 10c; emission plot, compare CS2 and CS3). However, the association of middle-S with active histone marks appears contradictory to our genome-wide correlation results (Figure 5). We attribute this apparent discrepancy to the fact that the CS3 is only a small proportion (∼4%, Supplementary Figure 10a, CS3) of the total genome, whereas all of middle-S is nearly one third of the genome. The active gene chromatin histone acetylation mark H3K27ac had correlation values of *r* = +0.72 for early-S versus r = –0.61 for middle-S chromatin. Similarly, but in the opposite direction, the gene-repressive chromatin mark H3K9me2 had correlation values of *r* = –0.19 for early-S versus r = +0.044 for middle-S chromatin (Figure 5) consistent with previous work ^2^. In addition, transcriptional characteristics of genomic regions, including transcription start site mapping and transcriptomics, also tend to mirror the pattern of epigenomic marks with regard to gene activity. Specifically, analyzing expression levels for promoters using the RAMPAGE assay ^66,79^, we observed a correlation of *r* = +0.55 for early-S chromatin and *r* = –0.12 for middle-S chromatin.

Both chromatin state and genome-wide correlation analyses indicate that middle-S shares more characteristics with early-S than it does with late-S. This result is consistent with the replicative labeling patterns that initially led us to classify middle-S as euchromatin. However, chromatin state analysis with RT inclusion also revealed that some early-S regions (CS1, 14% of the genome) as well as some middle-S regions (CS4, 18% of the genome) lack the active epigenomic mark associations and have a high overlap with transposable elements (Supplementary Figure 10e). These findings further illustrate the fact that TEs in the maize genome can populate regions considered by multiple criteria to be euchromatic.

The FISH experiments also provided new and compelling cytological evidence for the bifurcation of the global A compartment into early and middle chromatin. Unlike our previous work, which was limited to replicative labeling within early-S or middle-S, our RT-FISH approach allowed us to examine the spatial location of the RT-defined sequences outside of S phase and independent of replicative labeling. We carried out two related types of experiments. The first experiment used early and middle RT FISH probes separately on total nuclei and quantified the DAPI concentration in the FISH-defined regions relative to a similarly sized chromatin control region with properties typical of bulk euchromatin (Figure 6). The second experiment combined early and middle probe sets labelled with different fluorophores, hybridizing them together on purified G1 nuclei. This procedure enabled us to simultaneously visualize the spatial separation and nuclear localization of both RT-FISH signals, as well as to quantify their relative DAPI densities, without the need for an external control region (Figure 7). In every case, we observed signal partitioning that validates the two-compartment architecture, which we now know is present in G1 nuclei. This finding demonstrates that RT-related nuclear architecture exists prior to S phase, and is not simply a result of processes in S phase itself.

Given that Hi-C compartments and RT are tightly coupled in root tip, we reasoned that one could ask to what extent RT programs might be conserved in other tissues, using Hi-C structure as a proxy for RT data. We showed that biologically vastly different organs, earshoot versus root tip, had strong correlations for Hi-C based compartmentalization genome-wide, with *r* = +0.85 for both EV and IS (Supplementary Figure 5). When correlating RT from root tip with Hi-C measures from earshoot or bundle sheath, the correlations were similar in strength and direction to those from within-tissue comparisons (Figure 3). These findings provide evidence in support of the idea that the two-compartment architecture of euchromatin (as depicted in Figure 1f-h) not only extends throughout the cell cycle in root tips but appears to exist in other tissues as well. This observation raises the intriguing possibility that this pattern of nuclear architecture may reflect a broader organizational principle in other large-genome eukaryotes.

## AUTHOR CONTRIBUTIONS

HWB, LH-B and WFT conceived the study. HSA, HWB, LC, LH-B and WFT designed the experiments. HSA prepared Hi-C libraries and performed 3D FISH experiments, collected and analyzed the data. ZMT prepared MMOA-seq libraries. LC and HWB contributed to the analysis and interpretation of data. EEW and LM-Y provided maize root tip nuclei for the study. HSA and HWB drafted the manuscript and all authors critically revised and approved the final manuscript.

## COMPETING INTERESTS

We do not have any competing interests.

## MATERIAL AND CORRESPONDENCE

Material request from the corresponding author Hank W. Bass.

## ACKNOWLEDGEMENTS

The authors would like to acknowledge help with early versions of the data analysis by Dr. Jawon Song and Joshua Urrutia at the Texas Advanced Computing Center. This work was supported by a grant from the National Science Foundation Plant Genome Research Program (NSF IOS 2025811 to LH-B, LC, WFT, and HWB), awards from Florida State University (Dean’s Award for Doctoral Excellence to HSA; Ben and Karen Thrower award to HSA), and funds from North Carolina State University.

## DATA AVAILABILITY

Raw Hi-C data, MMOA-seq data, processed all valid pairs files, Hi-C files and bigwig files can be accessed at NCBI GEO under the accession number GSE287128. Scripts for Hi-C data analysis can be accessed at https://github.com/Sarachaudry/HiC-contacts-with-RT-annotations https://github.com/Sarachaudry/Correlation_Hexbin_plots https://github.com/Sarachaudry/virtual-4C-interactions-with-Sushi-in-R https://github.com/Sarachaudry/ChromHMM-Chromatin-States-Analysis https://github.com/Sarachaudry/R-script-for-scatter-plot

GEO accession GSE287128

## SUPPORTING INFORMATION LEGENDS

**Supplementary Figure 1. Experimental flow for Hi-C library prep.**

**(a-c)** Workflow: **(a)** 3-day old seedlings, **(b)** excised 0-1 mm root tip for fixation and nuclei isolation, **(c)** nuclei stained with DAPI and analyzed using flow cytometry (DAPI fluorescence with emission filter 450 ± 40 nm). Total nuclei (black interval bracket) were collected for each biological replicate of Hi-C. **(d)** Overview of the Hi-C protocol: Cells are cross-linked with 1% paraformaldehyde, creating covalent links between spatially adjacent chromatin segments (DNA fragments and proteins mediating interactions). Chromatin is then digested with the restriction enzyme DpnII (10 U/µl). The resulting sticky ends are repaired with nucleotides, one of which is biotinylated (indicated by a purple dot). Ligation is performed under highly dilute conditions to favor intramolecular ligation events. The DNA is subsequently purified and sheared, and biotinylated junctions are isolated using streptavidin beads. Interacting fragments are identified through paired-end 150-bp sequencing.

**Supplementary Figure 2. SCC score and Hi-C contact matrix heatmaps.**

**(a)** SCC correlation score heatmap of all root tip bioreplicates. **(b)** Hi-C contact matrix heatmap of chromosome 5 with early, middle and late RT segments on the top and left of the matrix.

**Supplementary Figure 3. Correlation of RT with Hi-C eigenvector analyses at multiple resolutions.**

**(a)** Genome-wide correlation plots showing Repli-seq values (x-axis) versus Hi-C eigenvectors, for either chromosome-wide eigenvector (CEG) or block-based eigenvector (BEG) methods (y-axis), at multiple resolutions. Pearson correlation coefficients (r) are indicated in blue (early-S), green (middle-S), and red (late-S). Black lines represent fitted linear regression lines for early, middle or late data. **(b)** Bar graph of correlation coefficients (r) between Repli-seq and Hi-C eigenvectors across different bin sizes (x-axis). Correlations for early-, middle-, and late-S regions are shown as blue, green, and red bars, respectively. **(c)** UCSC Genome Browser view of chromosome 5 showing the Repli-seq profile and Repliscan segments with CEG (25, 50, and 100 kb resolution) and BEG (50kb resolution) eigenvector tracks below. The orange box highlights the 50-kb resolution used throughout the manuscript and the pink dashed box highlights the knob region **(d)** 2-Mb zoomed-in regions from a non-heterochromatic part of chromosome 5, illustrating finer-scale chromatin structure.

**Supplementary Figure 4. Correlation of RT with Hi-C insulation score analyses at multiple resolutions.**

**(a, b)** Bar graphs of genome-wide correlation coefficients (r) between Repli-seq and Hi-C insulation scores (IS) across different bin sizes (x-axis) with a sliding window size of either 2.5 Mb **(a)** or 1 Mb **(b)**. Correlations for early-, middle-, and late-S regions are shown as blue, green, and red bars, respectively. **(c)** UCSC Genome Browser view of chromosome 5 showing the Repli-seq profile and Repliscan segments with insulation score analyses at all resolutions and window sizes below. The orange box highlights the 50-kb resolution used for genomewide correlation analysis in the main text. **(d)** 2-Mb zoomed-in regions from a non-heterochromatic part of chromosome 5, illustrating finer-scale chromatin structure.

**Supplementary Figure 5. Nuclear architecture comparison among different tissues.**

**(a)** Genome-wide eigenvector (EV) analysis comparison between different tissues: root tip versus earshoot, root tip versus bundle sheath, and bundle sheath versus earshoot. **(b)** Genome-wide insulation score (IS) analysis comparison between the same tissue pairs.

**Supplementary Figure 6. Chromosome-wide comparison among different tissues.**

**(a)** Chromosome-wide correlation of chromatin structure between root tip (blue) and earshoot (red). **(b)** Chromosome-wide correlation of chromatin structure between root tip (blue) and bundle sheath (red). The Y-axis represents eigenvector values, while the X-axis denotes chromosome coordinates.

**Supplementary Figure 7. Contact matrix of other tissues.**

**(a)** A chromosome schematic with an orange rectangle highlighting a 2-Mb genomic region from the short arm of chromosome 5. **(b-c)** Hi-C contact matrix of the indicated 2-Mb region for earshoot tissue **(b)** or bundle sheath tissue **(c)**. The RT classification segments (early in blue, middle in green) are annotated along the top and left edges. Matrix color intensity represents the frequency of contacts between loci. Blue circles highlight examples of early-early contacts. **(d, e)** UCSC Genomaize browser views of the same 2-Mb region, displaying Repli-seq profiles (RT), eigenvector (EV) analysis, and insulation score (IS) tracks for earshoot (**d**) and bundle sheath (**e**). Blue and green shaded regions correspond to early and middle RT segments and their respective EV and IS patterns.

**Supplementary Figure 8. Distance normalization of Hi-C contacts.**

**(a)** Different RT contacts are normalized with respect to distance across the genome and plotted separately for early-early (blue), early-middle (gray), middle-middle (green), and late-late (red).

**(b)** Randomly shuffled RT class contacts with respect to distance (control).

**Supplementary Figure 9. Nuclear architecture versus chromatin marks.**

**(a-h)** Correlation plots across chromosome 5 comparing root tip Hi-C eigenvector (EV) analysis data (blue, on the right y-axis) with various chromatin marks, transcription and chromatin accessibility data (red, on the left y-axis) along chromosome 5 coordinates (x-axis) for **(a)** H3K27ac, **(b)** H3K4me3, **(c)** H3K4me1, **(d)** H3K56ac, **(e)** H3K27me3, **(f)** H3K9me2, **(g)** RAMPAGE transcription, **(h)** DNS-Light digest, and **(i)** MMOA-seq.

**Supplementary Figure 10. Summary of ChromHMM chromatin state models with and without Repli-seq integration.**

**(a)** Pie chart showing the percentage of each chromatin state in the maize genome for the 6-state model that includes Repli-seq (wRT) and eight epigenomic, transcriptional, and chromatin accessibility datasets. The 6 state map wRT shows a more balanced distribution of chromatin states. **(b)** Pie chart of chromatin state proportions using the same marks as in (a) but without the inclusion of Repli-seq data (woRT). The 6-state map woRT shows that CS1, CS4, CS5, and CS6 each cover only 3–5% of the genome. In contrast, CS2 and CS3 are the most abundant states. **(c)** ChromHMM emission and transition probability heatmaps for the wRT model; color intensity represents the level of mark enrichment/probability. **(d)** Base-pair overlap between Repli-seq defined RT classes and the chromatin states wRT model( blue for early-S, cyan for early-middle, green for middle-S, yellow for middle-late, red for late-S, and grey for others (e.g., early-late, early-middle-late, or no RT). **(e)** Base-pair overlap of gene and transposable element (TE) annotations with the chromatin states wRT model. Light green indicates gene overlap; light blue indicates TE overlap. **(f)** ChromHMM emission and transition probability heatmaps for the woRT model; color intensity indicates mark enrichment/probability. **(g)** Base-pair overlap between Repli-seq RT classes and chromatin states woRT model. **(h)** Base-pair overlap of genes and TEs with chromatin states woRT model. (i) Sum of off-diagonal transition probabilities in ChromHMM models trained with 3 to 12 chromatin states, comparing models with RT (light grey) and without RT (dark grey) included. This allows for extended comparison of different models.

**Supplementary Figure 11. Browser view of chromatin states.**

**(a)** Genome browser view comparing two different chromatin state model types in selected regions on maize chromosome 5. Two genes are highlighted in dark grey vertical bars with their flanking regions in light grey. Tracks shown include Repli-seq profiles and RT segments, TE and gene annotations, chromatin states (CS) from the with RT model, ChIP-seq, RAMPAGE transcription, DNS-light digest and chromatin states from the without RT model. **(b)** Zoomed-in views of the highlighted gene regions from panel (a). Interestingly, the inclusion of Repli-seq data in chromatin state modeling tends to capture genic and flanking regions into singular states CS2 and CS1 respectively, compared to the more fragmented multi-state pattern seen for these same genes in the models trained without Repli-seq

**Supplementary Figure 12. ChromHMM emission probability heatmaps.**

Emission probability heatmaps of ChromHMM models trained with RT data in 3-, 4-, 5-, and 6-state configurations. The color intensity represents the enrichment and probability of each input dataset.

**Supplementary Figure 13. FISH probes designed for chromosome 5 short arm.**

**(a)** Percentages of early and middle RT segments are shown for the whole maize genome, chromosome 5, and the short arm of chromosome 5 (5S). The 5S region of chromosome 5 was selected as a representative genomic region for 3D FISH analysis due to its similar RT segment distribution. **(b)** Table displaying the total number of early and middle FISH probes and their density.

**Supplementary Figure 14. FISH probe labeling.**

**(a)** Workflow for designing fluorescent probes for 3D FISH experiments. A 45-mer oligonucleotide library was constructed with common end-flanking sequences for the attachment of forward and reverse primers. Libraries for early and middle RT probes were amplified and transcribed to produce single-stranded RNA copies using T7 in vitro transcription. The resulting RNA was reverse transcribed into cDNA using an RNA-DNA chimeric primer. Various fluorescent dyes (AF-488, AF-647, and AF-548) were introduced at this stage. The RNA was then hydrolyzed using RNase-A. The resulting single-stranded DNA (ssDNA) labeled with different colored fluorophores was used for 3D FISH experiments. **(b-d)** Experimental workflow: **(b)** 3-day old seedlings were subjected to EdU labeling for 20 minutes. **(c)** Immediately after EdU labeling, the terminal 0–1 mm root tip was excised, fixed, and root tip nuclei were isolated. **(d)** EdU was conjugated to a fluorophore (AF-488) using click chemistry and the nuclei were counterstained with DAPI. Nuclei were analyzed via flow cytometry using 355 nm (UV) and 488 nm (blue) lasers. The bivariate plot displays DNA content (DAPI fluorescence) and EdU incorporation (AF-488 fluorescence). Rectangles indicate the G1, early S, middle S, late S and G2 nuclei that were sorted and then combined together for the entire cell cycle mixed nuclei pool shown in Figure 6. Sorted G1 nuclei were used for FISH experiments shown in Figure 7. **Supplementary Table1: Hi-C summary statistics**

## Notes

### Competing Interest Statement

The authors have declared no competing interest.

