## Supplemental Figures and Table for "Replication Timing Uncovers a Two-Compartment Nuclear Architecture of Interphase Euchromatin in Maize"

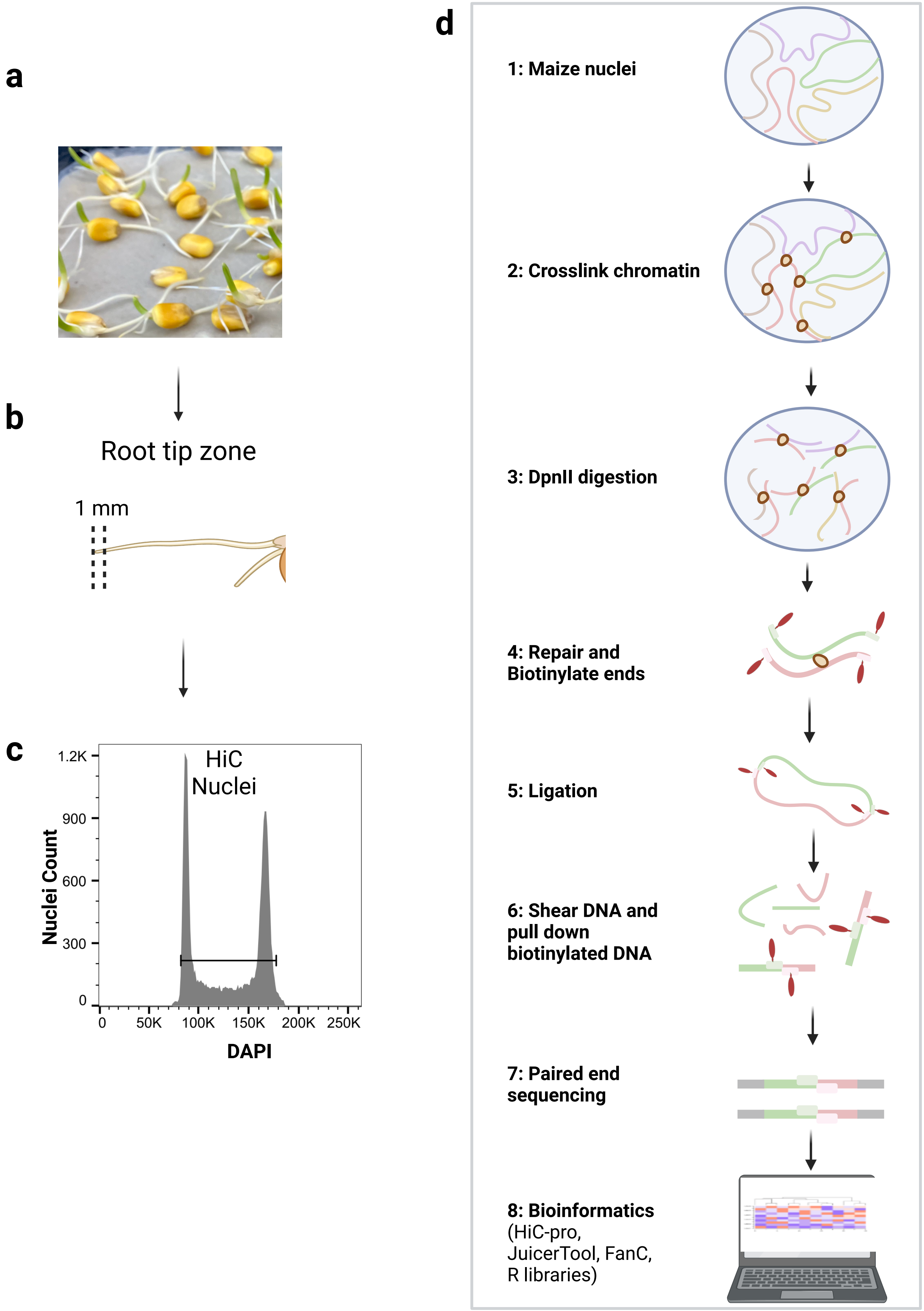


Akram, S Figure 1

**a**

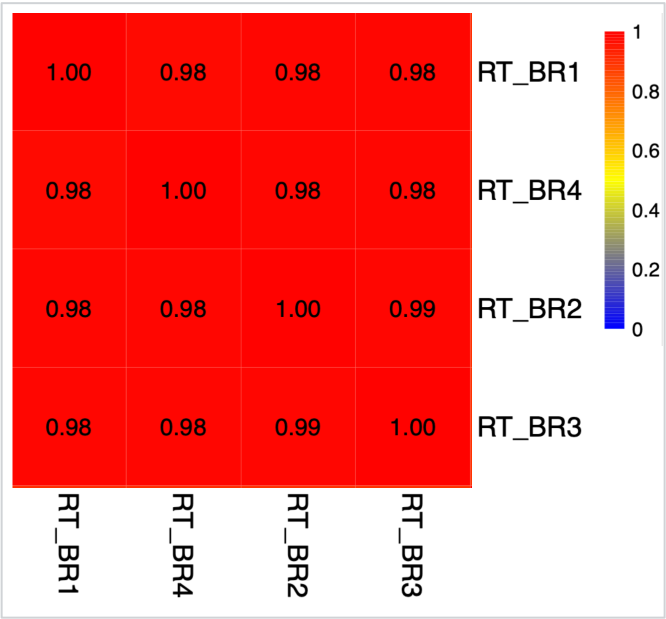


**b**

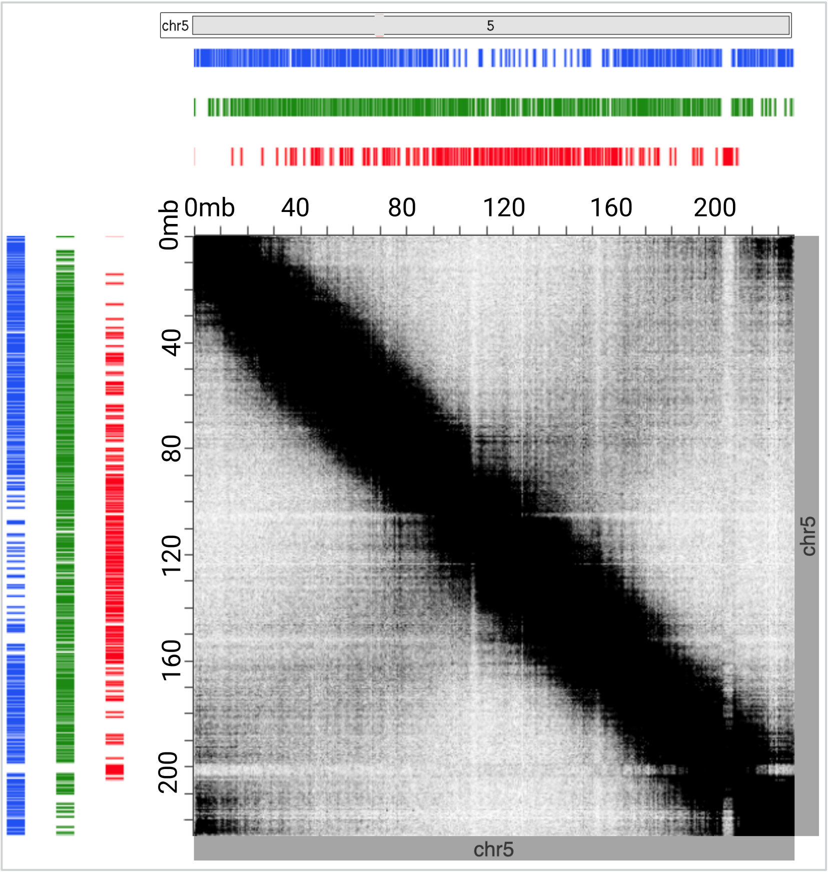


Akram, S Figure 2

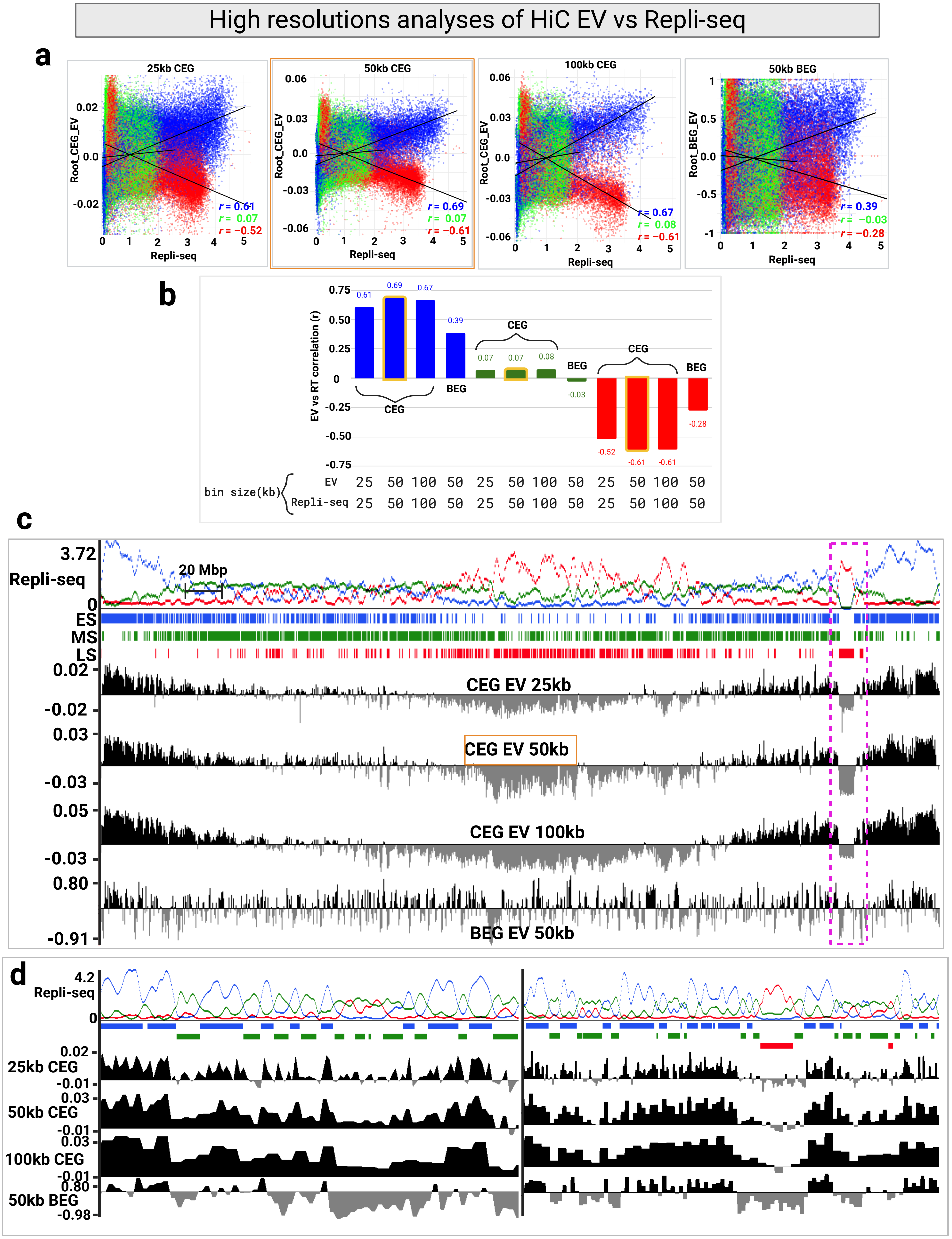


Akram, S Figure 3

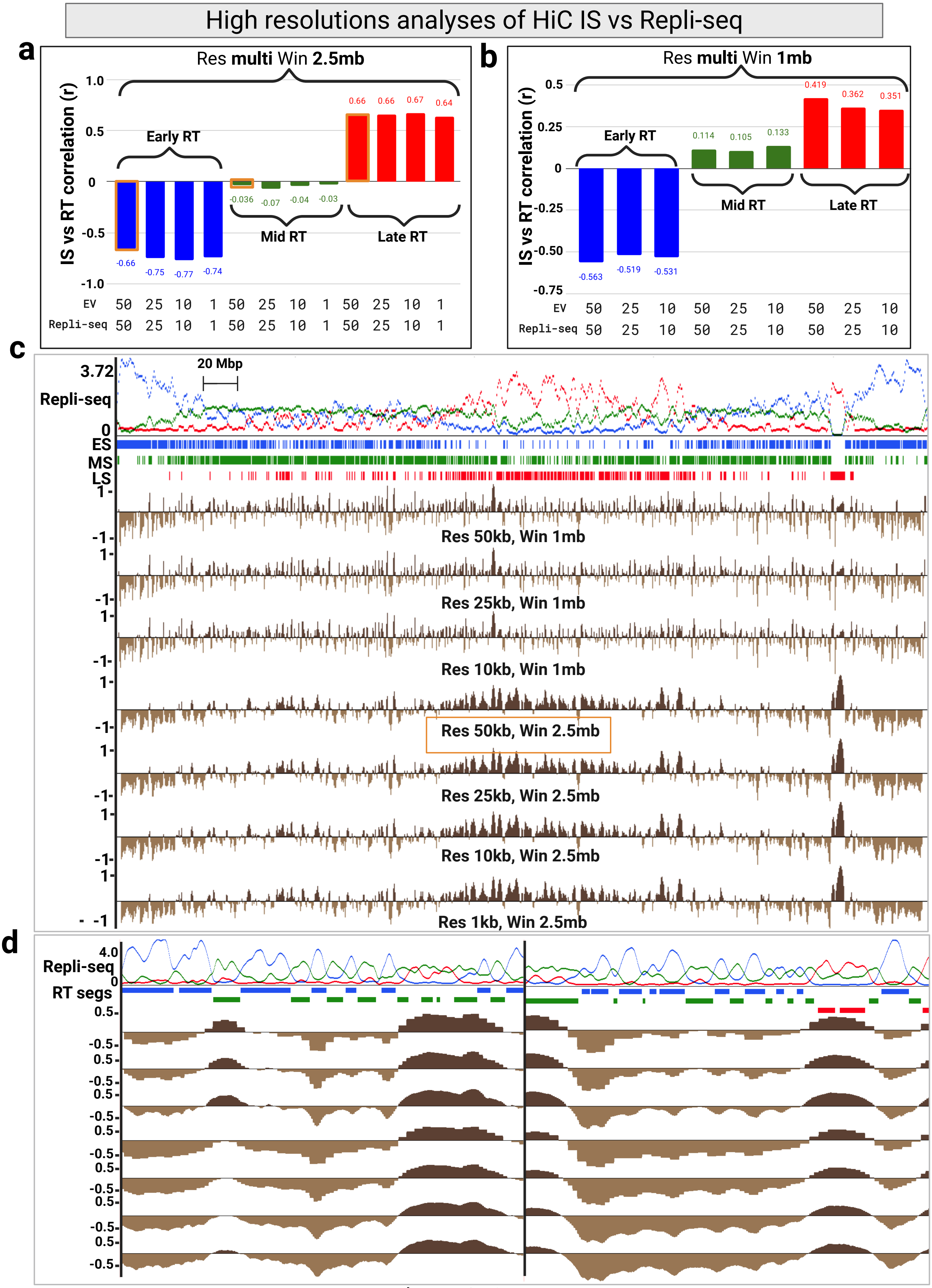


Akram, S Figure 4

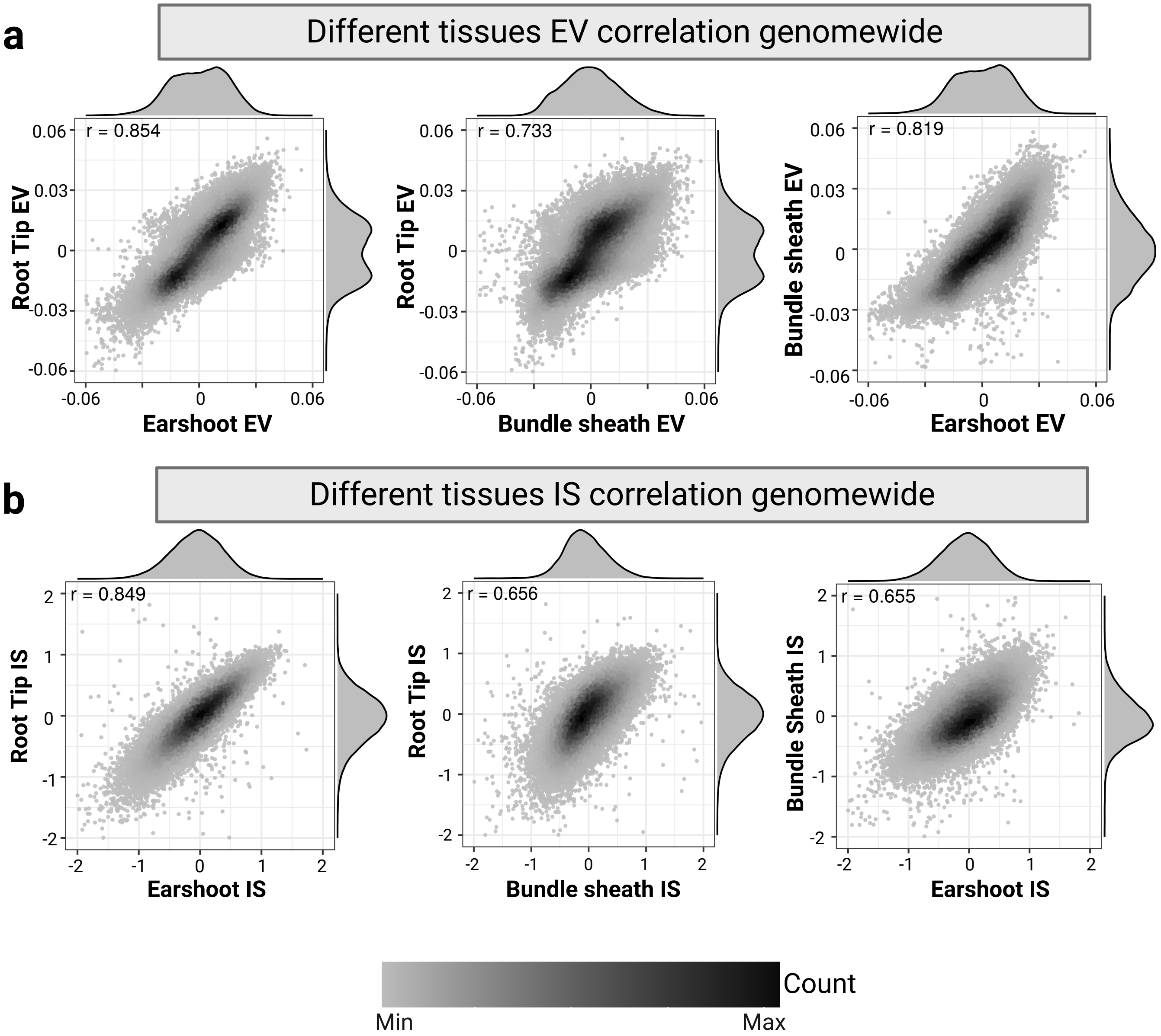


Akram, S Figure 5

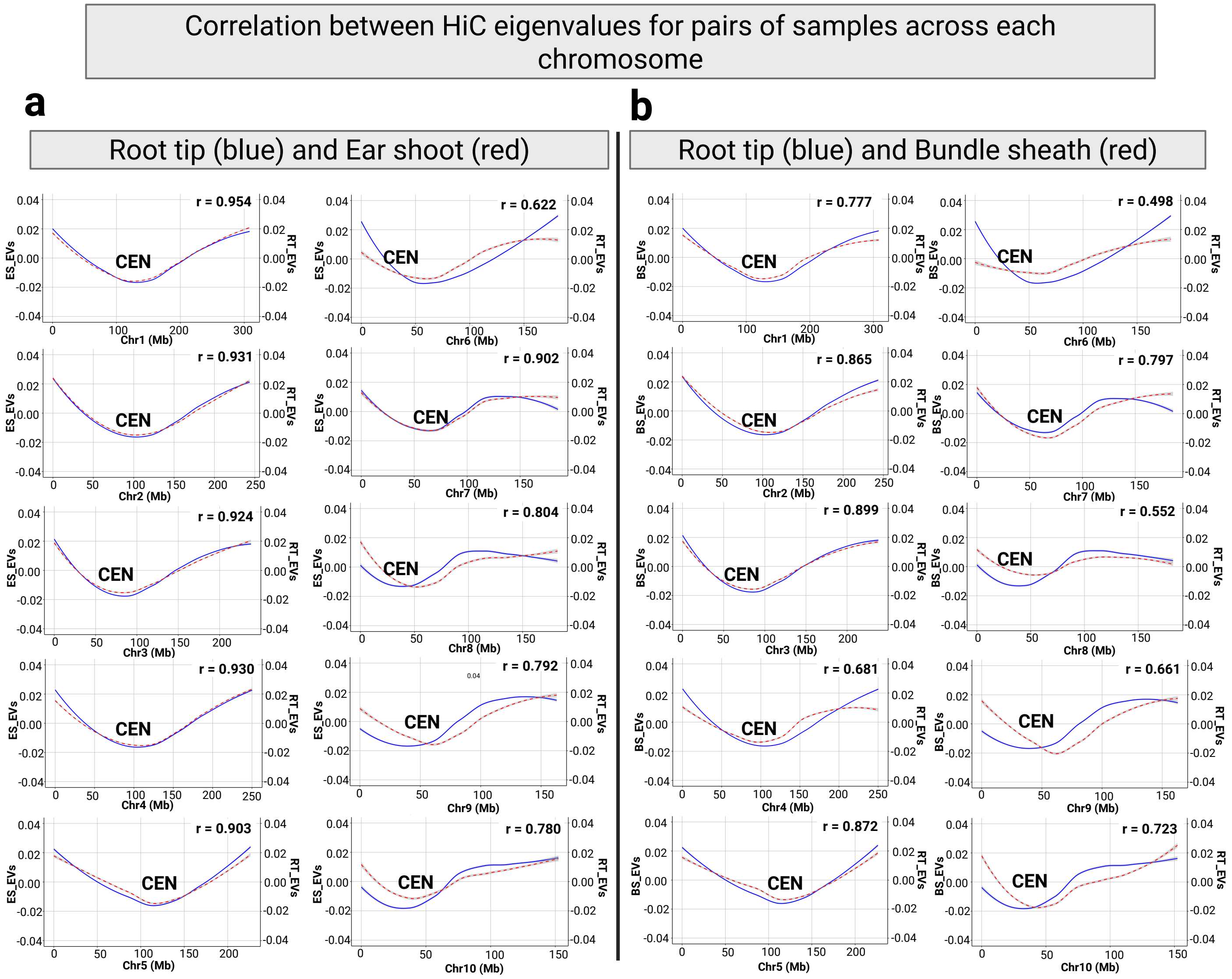


Akram, S Figure 6

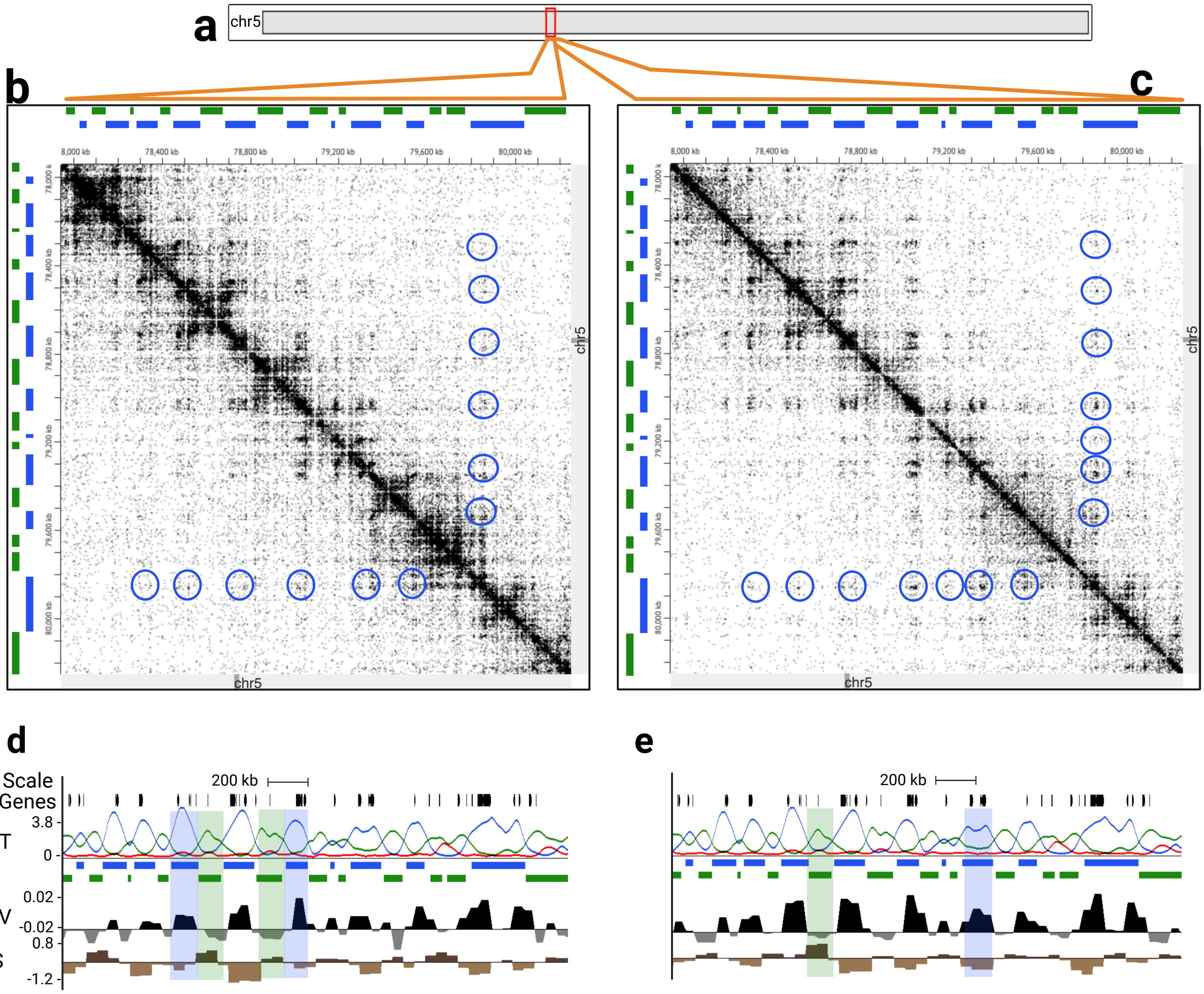


Akram, S Figure 7

### EvsE and MvsM contact pair frequencies normalised by distance

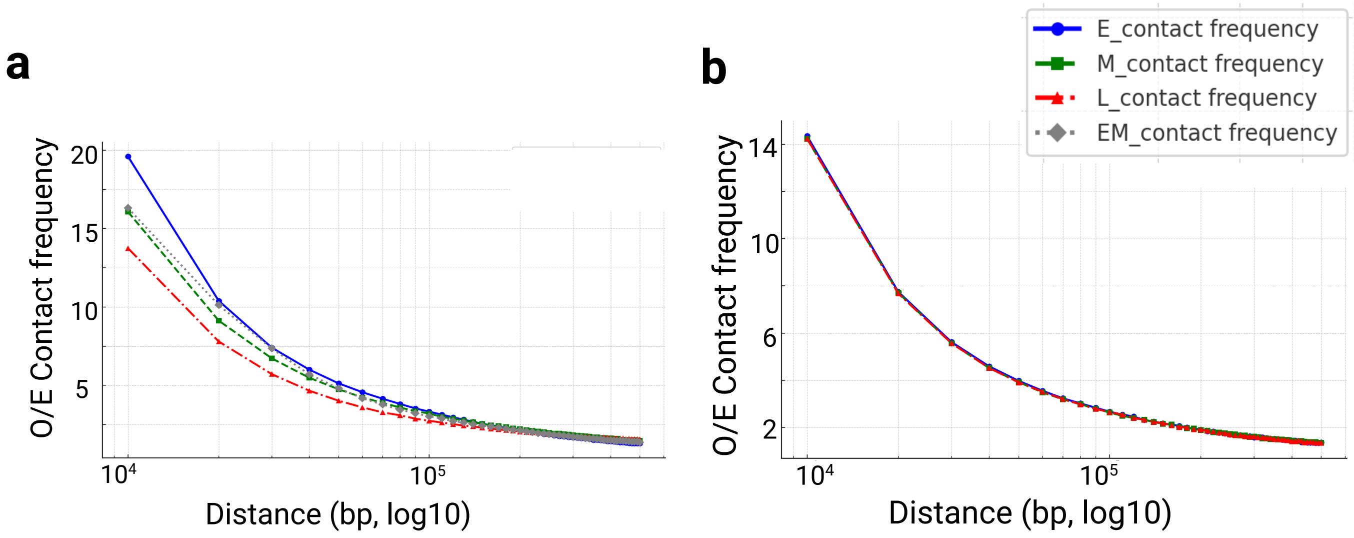


Akram, S Figure 8

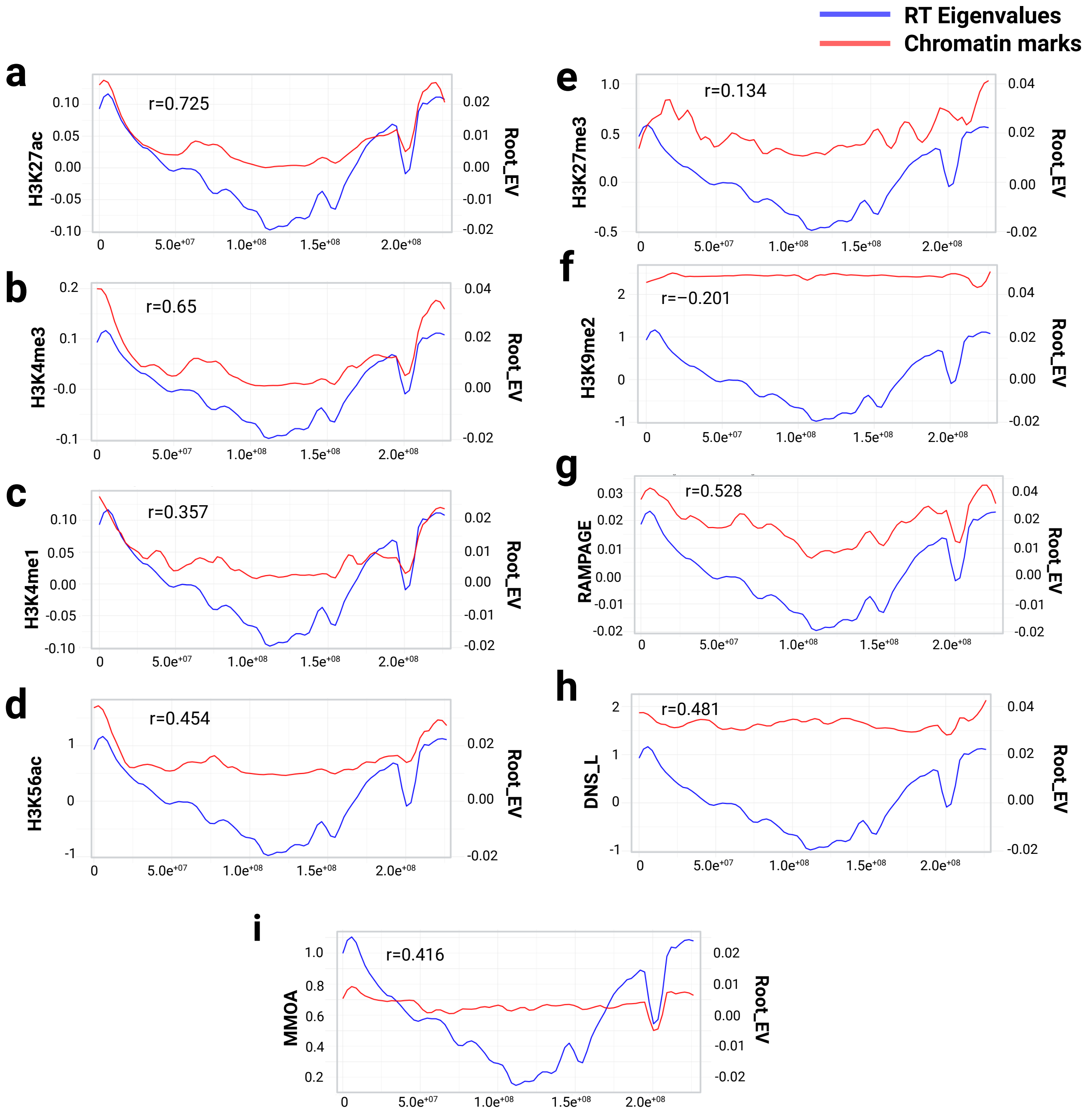


Akram, S Figure 9

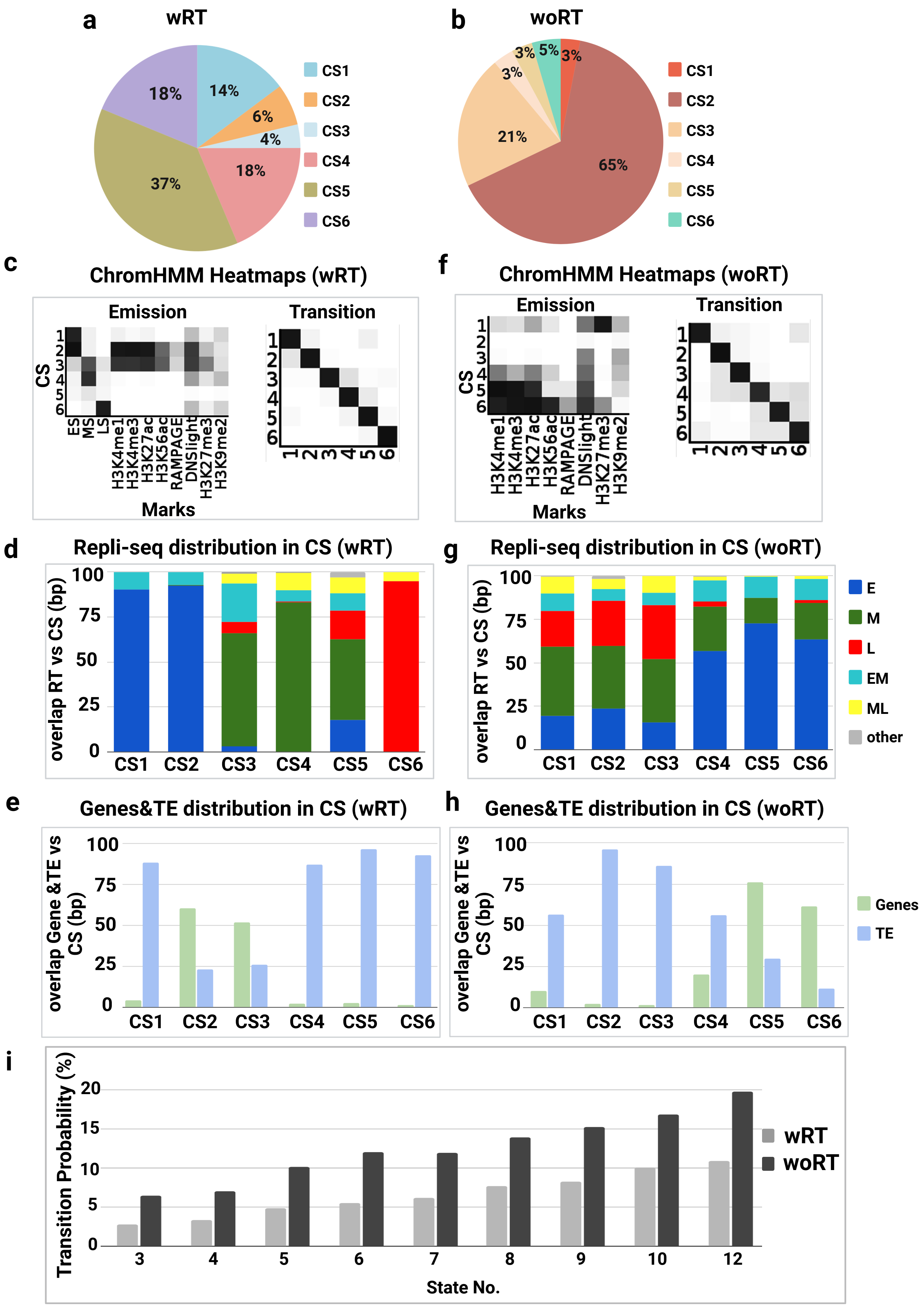


Akram, S Figure 10

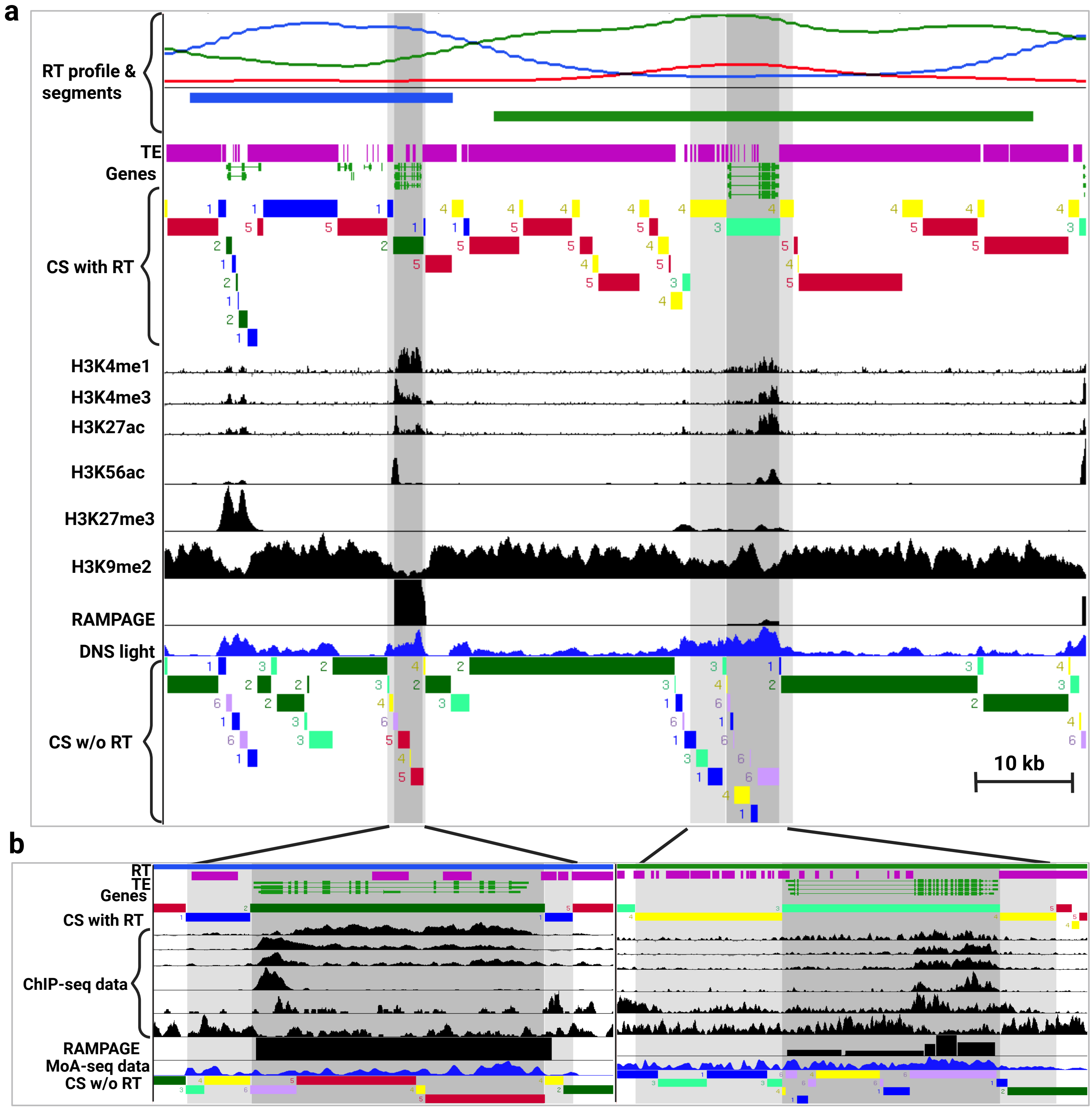


Akram, S Figure 11

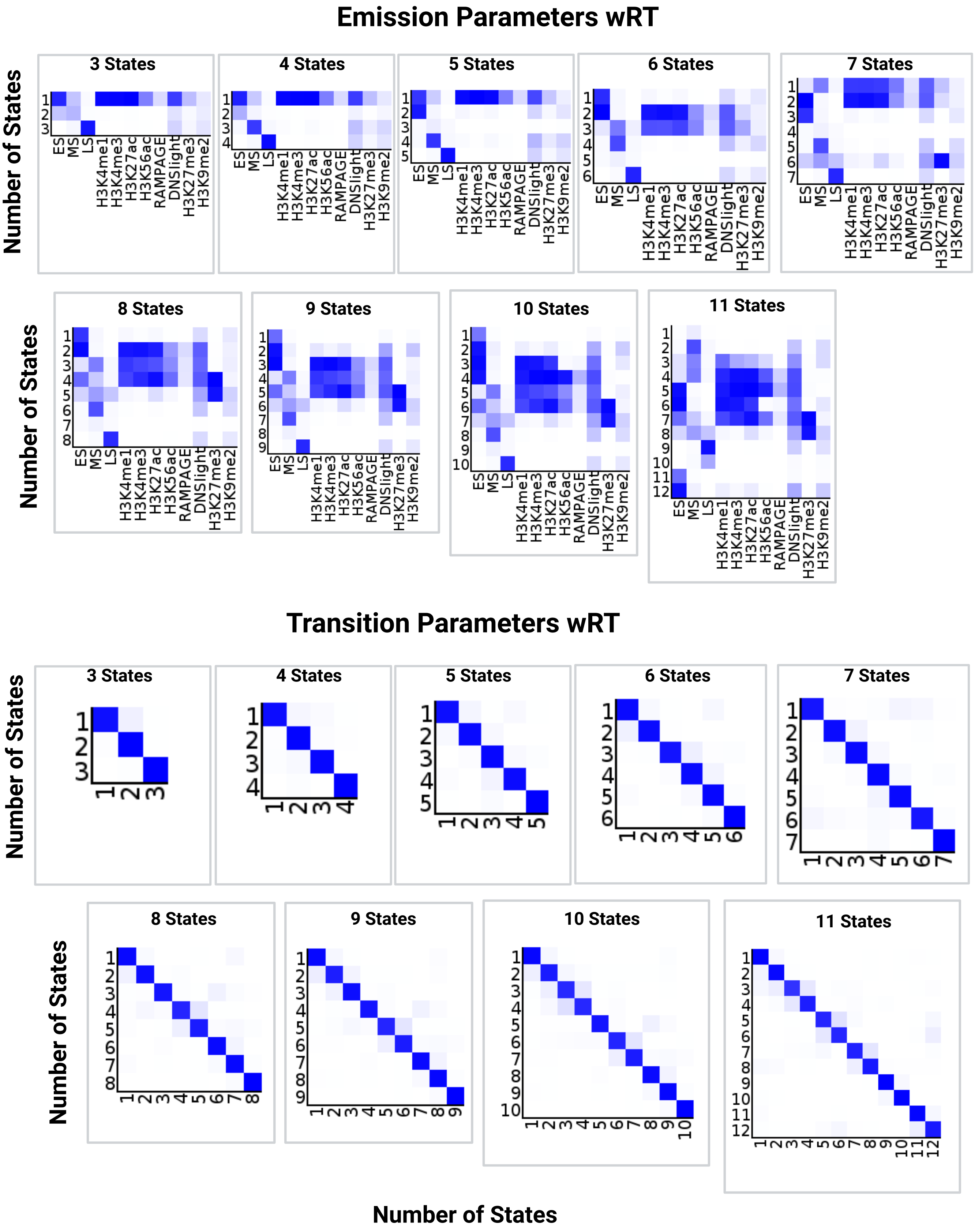


Akram, S Figure 12

a

### Genomic region for FISH

| Genomic region | Number of segments (bp, % whole genome) | |
|---|---|---|
| | EARLY | MIDDLE |
| Chromosomes 1-10 (2180 MB) | 4407 (560MB, 30%) | 7641 (730MB, 35%) |
| Chromosomes 5 (226 MB) | 486 (69MB, 30%) | 797 (77MB, 34%) |
| 5S, short arm of 5 (105 MB) | 270 (34MB, 30%) | 427 (42MB, 38%) |

b

### FISH probes for chr5 short arm

| RT Segments | Probe number | Probe density |
|---|---|---|
| Early | 22,460 | 0.66 / kb |
| Middle | 27,390 | 0.65 / kb |

Akram, Figure 13

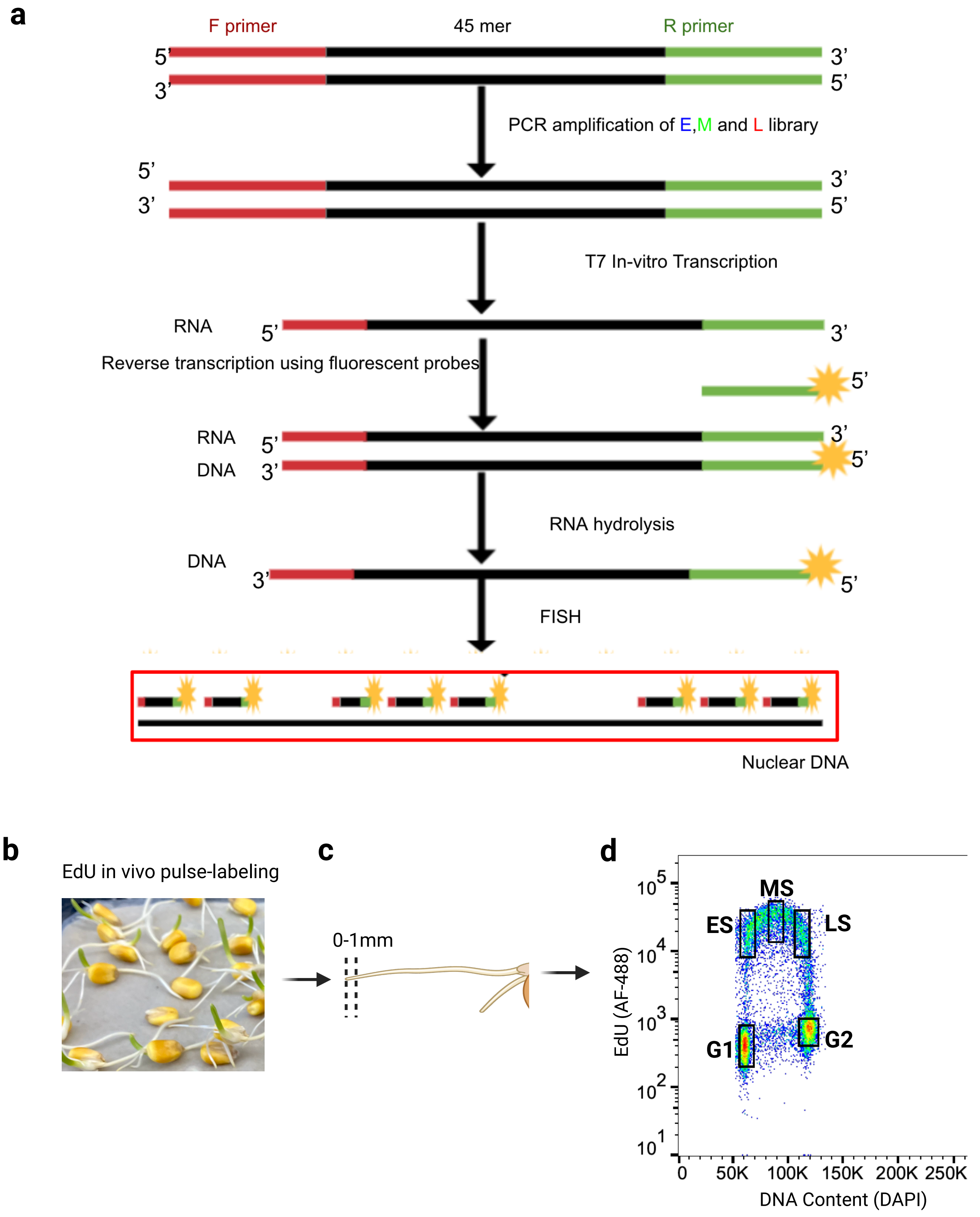


Akram, S Figure 14

Supplementary Table 1. HiC Summary Statistics.

| HiC libraries[A] | Total Reads[B] | Valid Contacts[C] | PCR Duplicates | Trans Interactions | Cis Interactions | Cis Short Range | Cis Long Range |
|---|---|---|---|---|---|---|---|
| **Root tip rep1** | 148,529,729 | 46,277,361 | 8,646,492 | 5,815,255 | 31,815,614 | 6,593,800 | 25,221,814 |
| **Root tip rep2** | 146,330,218 | 42,912,077 | 8,065,453 | 7,005,296 | 27,841,328 | 5,889,328 | 21,952,000 |
| **Root tip rep3** | 187,537,189 | 58,057,354 | 9,410,986 | 9,507,348 | 39,139,020 | 7,684,329 | 31,454,691 |
| **Root tip rep4** | 143,438,961 | 41,549,774 | 6,834,567 | 7,190,483 | 27,524,724 | 5,603,289 | 21,921,435 |
| **Ear shoot rep1** | 127,467,879 | 27,730,693 | 4,505,474 | 4,769,700 | 18,455,519 | 4,162,271 | 14,293,248 |
| **Ear shoot rep2** | 308,506,330 | 95,879,933 | 18,220,265 | 20,675,903 | 56,983,765 | 8,711,364 | 48,272,401 |
| **Ear shoot rep3** | 253,912,920 | 74,369,477 | 14,472,034 | 17,325,131 | 42,572,312 | 42,572,312 | 35,915,439 |
| **Bundle sheath rep1** | 448,399,207 | 110,363,506 | 39,723,571 | 36,158,855 | 74,204,651 | 18,148,355 | 56,056,296 |
| **Bundle sheath rep2** | 867,566,517 | 294,700,726 | 86,109,501 | 88,319,269 | 206,381,451 | 51,183,865 | 155,197,592 |

Footnotes:

A. Root tip and Ear shoot samples from this study. Bundle sheath libraries are from Dong *et al.,* (2017)[32].
B. Total read pairs are from initial sequencing.
C. The remaining columns are output values from HiC-pro pipeline Servant *et al*., (2015)[33].
